# Automated Prediction of Fibroblast Phenotypes Using Mathematical Descriptors of Cellular Features

**DOI:** 10.1101/2024.05.15.594418

**Authors:** Alex Khang, Abigail Barmore, Georgios Tseropoulos, Kaustav Bera, Dilara Batan, Kristi S. Anseth

## Abstract

Fibrosis is caused by pathological activation of resident fibroblasts to myofibroblasts that leads to aberrant tissue stiffening and diminished function of affected organs with limited pharmacological interventions. Despite the prevalence of myofibroblasts in fibrotic tissue, existing methods to grade fibroblast phenotypes are typically subjective and qualitative, yet important for screening of new therapeutics. Here, we develop mathematical descriptors of cell morphology and intracellular structures to identify quantitative and interpretable cell features that capture the fibroblast-to-myofibroblast phenotypic transition in immunostained images. We train and validate models on features extracted from over 2,500 primary heart valve interstitial cells (VICs) and test their predictive performance on VICs treated with the small molecule drug 5-azacytidine, which inhibited myofibroblast activation. Collectively, this work introduces an analytical framework that unveils key features associated with distinct fibroblast phenotypes via quantitative image analysis and is broadly applicable for high-throughput screening assays of candidate treatments for fibrotic diseases.

## INTRODUCTION

Fibrosis can occur in virtually any organ including the heart, kidney, liver, skin, and lungs and is implicated in nearly 45% of deaths in the modern world^1^. Moreover, fibrosis and cancer associated fibroblasts are known to pay a role in tumor metastasis^2^. Fibrosis occurs when normal wound healing processes are dysregulated leading to excessive deposition of structural extracellular matrix (ECM) components, such as collagen, and results in aberrant stiffening of tissues and organs. If left unresolved, the fibrotic process causes abnormal remodeling of the ECM, organ malfunction or even failure, and in severe cases, death^3^. To date, only two drugs, OFEV^®^ (nintedanib) and Esbriet^®^ (pirfenidone), are approved to treat fibrosis but are marketed only for the treatment of idiopathic pulmonary fibrosis (IPF)^4^.

While the role of the fibroblast to myofibroblast transition is well recognized in fibrosis and disease progression, the complex nature of the disease, especially its initiation and progression as in the case of IPF^5^, has led to a scarcity of pharmaceutical therapies. Typically, fibroblasts reside within stromal tissue and are responsible for the secretion and remodeling of ECM components, thereby playing a critical role in many wound healing processes. In response to insult, quiescent fibroblasts undergo a phenotypic activation to myofibroblasts characterized by increased proliferation, ECM synthesis, and contractility mediated by expression of alpha-smooth muscle actin (αSMA) stress fibers. When periods of repeated or severe insult occur, the tissue-resident myofibroblast phenotype can persist, which results in excessive matrix deposition and pathological tissue/organ stiffening. Subsequently, excessive tissue/organ stiffening further encourages myofibroblast differentiation and initiates a positive feedback loop. Due to their pivotal role in fibrosis, fibroblasts and their activation to myofibroblasts have been extensively studied in virtually every organ system^6–9^.

A hallmark and widely used indicator of myofibroblast activation is the presence of discrete αSMA stress fibers^10^. Traditionally, researchers and pathologists have used immunostaining of tissue sections or fibroblast cultures to visualize αSMA; however, grading the severity and discreteness of the fibers has remained largely subjective. Prior published methods have used absolute or normalized fluorescence intensity of αSMA+ cells or tissues, but these measurements can vary with sources of antibodies, staining protocols, and different microscopes and imaging parameters (e.g., laser power and gain). Moreover, significant information, such as cell morphology, goes largely unused in fibroblast phenotyping. The few studies that have considered cell morphological measurements used qualitive assessments to classify cells^11^. Despite the widespread occurrence of fibrotic diseases and the desire to discover drugs that can target pathological myofibroblasts, a universally accepted method for quantifying fibroblast phenotypes is currently lacking and further complicates cross-study comparisons.

Towards automated and objective methods to characterize fibroblast phenotypes, researchers have begun to use neural networks to differentiate between quiescent and activated fibroblasts based on manually curated training data^12^. Although effective, these models require substantial training datasets, and the predictions are typically based on abstract features with limited physical and/or physiological interpretations. Furthermore, neural networks often exhibit high complexity, which might be unnecessary for the categorical classification of fibroblasts. As an alternative approach, others have used cell and nuclear morphologies to classify fibroblasts, which provides an avenue to better understand *how* myofibroblasts are different and not simply *if* they are different^13–15^. However, to date, no approach exists to combine morphological features and αSMA stress fiber architecture to grade fibroblast phenotypes.

In the present study, we developed a quantitative image analysis approach that includes both cell morphological features and expression of discrete αSMA stress fibers to demarcate quiescent and activated fibroblast phenotypes and to visualize their shapes across the activation spectrum. Our efforts focus on these markers and measurements, as they are the most widely used characteristics to delineate fibroblast phenotypes^6,11– 13,16,17^. As a result, data sets of fluorescence images are routinely collected to characterize fibroblast and myofibroblast populations and are readily available. Motivated by the prevalence of valve and cardiac fibrosis in the general population^18,19^, we use valve interstitial cells (VICs) isolated from porcine aortic valves and cardiac fibroblasts (CFs) isolated from adult rat ventricles as model fibroblast cells. VICs and CFs were induced into an activated state using both mechanical (soft vs stiff microenvironment) and biochemical (TGF-β) stimuli, respectively, to test and validate our analysis approach on several scenarios that promote myofibroblast activation. First, we train and validate models on the VIC dataset and then use the model to grade a test data set comprised of VICs seeded on stiff substrates and treated with the small molecule drug 5-azacytidine, which inhibits myofibroblast activation. As a final example, we extend the VIC-trained models to grade the CF dataset and compare the grading to models trained directly on the CF dataset (Appendix A). In our image analysis methodology, we use Fourier series to mathematically describe segmented fibroblast shapes and use the variable coherence to quantify the anisotropy of αSMA stress fibers in a manner independent from absolute fluorescence intensity. This information is then combined and used in exploratory data analysis to discover, and importantly *visualize*, the greatest variation between fibroblasts and myofibroblasts. Physical interpretation of these differences allows for the identification of morphological features (e.g., cell area, aspect ratio) that vary significantly between quiescent and activated fibroblasts. Finally, we test the ability of minimal key features to predict fibroblast phenotype and compare the automated algorithm to manual grading done via an independent visual assessment. Overall, we report a method using mathematical descriptors of cell morphological features to discover key measures that can predict fibroblast phenotypes with high levels of accuracy using a straightforward, but robust, method. Looking forward, the developed methodology can minimize human bias and serve as a high throughput method for grading fibroblast phenotypes during drug screening. Moreover, we anticipate that the established methodology may be extended to analyze fibroblast phenotypes in situ and augment histological assessment of diseased tissues. In a broader context, the current method presents the potential of a gold standard across research groups to ensure consistent fibroblast grading that would largely bolster the field of fibrosis research and expedite drug discovery.

## RESULTS

### Mathematical descriptors of cell and nuclear shapes

Cell and nuclear shapes are known to be influenced by changes in gene and protein expression, and in this regard, can be viewed as proxies of information regarding phenotypic state^14,15,20–22^, rendering methods to quantify cell morphologies useful towards understanding cell function. Here, we extended Fourier analysis^23–25^ to create mathematical descriptions of cellular (Fig. 1a) and nuclear (Fig. 1b) shapes and produce faithful reconstructions of their boundaries (Fig. 1c) from an optimized number of coefficients. Fourier analysis enables the reconstruction of arbitrary 2D shapes with varying degrees of precision, determined by the user-defined number of Fourier harmonics (*n)* (Fig 1d). Two parametric functions *x(θ)* and *y(θ)* are employed to relate the arc length around a given cell/nuclei boundary *(θ)* to the corresponding *x-* and *y-* Cartesian coordinates. Here, the arc length of the shape is normalized such that the total length is 2*π*. Essentially, this results in a one-to-one mapping of the original shape to a unit circle, ensuring that every cartesian *x,y* point corresponds to exactly one arc length *θ* in the range [0, 2*π*]. The equations *x(θ)* and *y(θ)* take the form

**Fig. 1:**
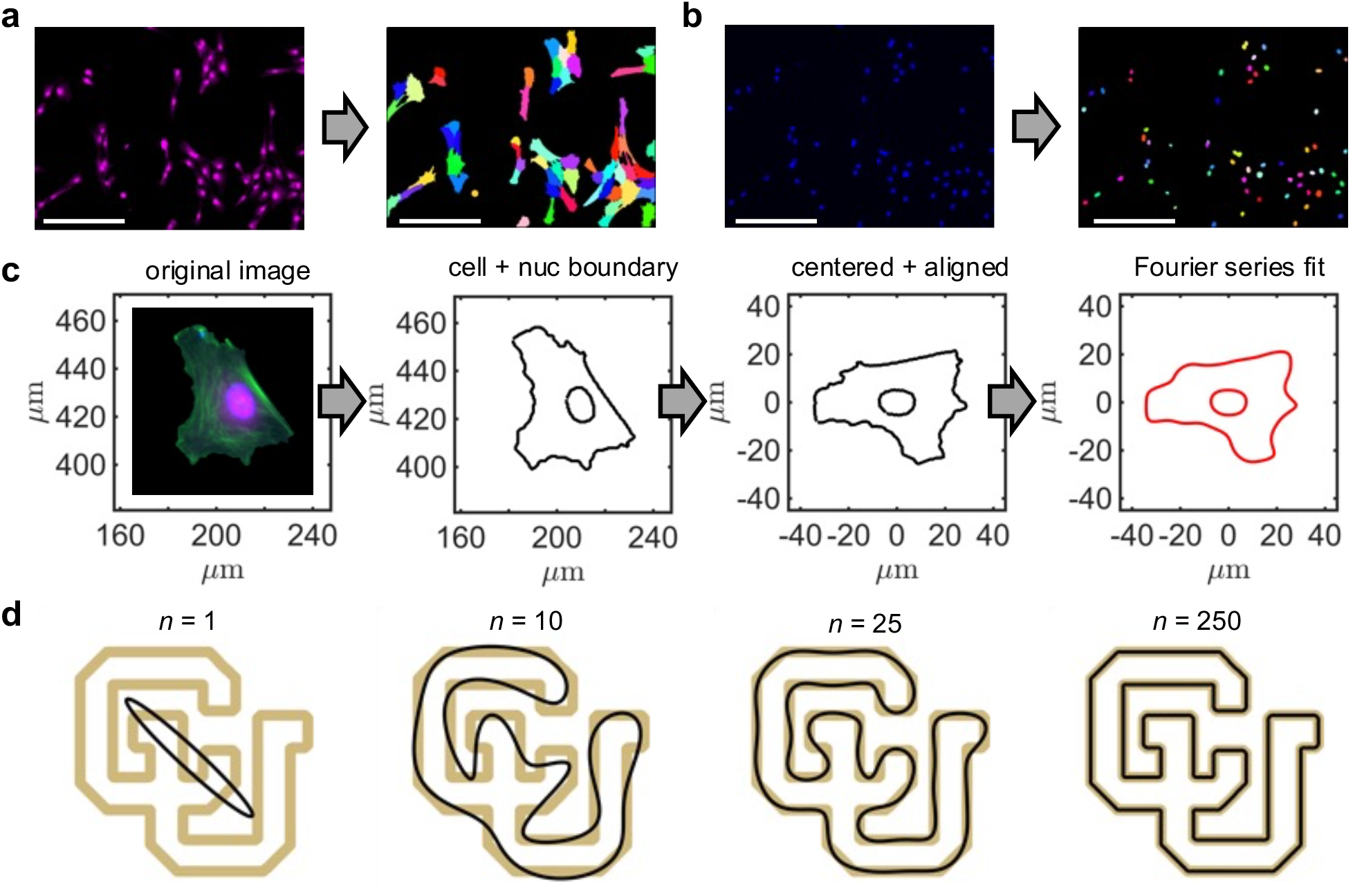
Segmentation and mathematical representation of VIC body and nuclear morphologies. **a**, The VIC cytoplasm is imaged and segmented using a marker-based watershed algorithm with the **b**, segmented nuclear geometries serving as markers. **c**, VIC morphological analysis is achieved through segmenting the cell and nuclear boundaries, centering, alignment of the VIC along its longest axis, and fitting a Fourier series expansion to the VIC body and nuclear shapes (body: *n* = 15, nuclei: *n* = 5). **d**, Fourier series expansion used to represent the geometry of a closed shape. Note that as the harmonic number *n* increases, the Fourier series expansion (black outline) converges to the target shape (gold outline). Stained images: green - αSMA, magenta – cytoplasm, blue – nuclei. Scale bars: 200 *µ*m.

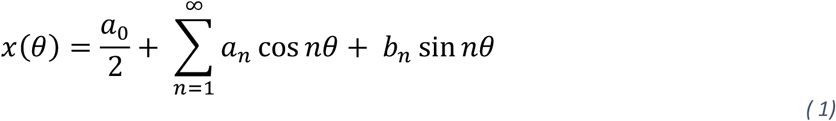

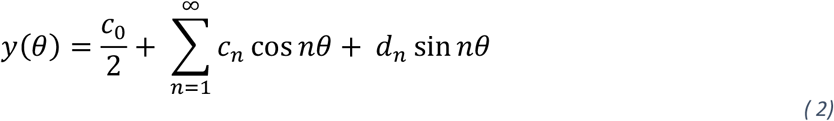

where *a*_*0*_, *a*_*n*_, *b*_*n*_, *c*_*0*_, *c*_*n*_, and *d*_*n*_ are Fourier coefficients determined by solving overdetermined systems of linear equations. Ultimately, this process yields Fourier coefficients that contain information regarding cellular morphology. However, it is important to note that the physical meaning of the coefficients is not obvious and instead offers a unique opportunity to employ a data-driven approach to analyze cell shapes. Here, the Fourier series representations of VIC body and nuclear geometries reduces the total number of variables being analyzed by representing many boundary points with appreciably fewer coefficients (e.g., >1,000 boundary points are reproduced accurately with 50 Fourier coefficients). Furthermore, the Fourier series produces a smoothed representation of the target shape when *n* is optimized and is key to recovering naturally smooth cell body and nuclear boundaries that can be distorted when represented by discrete pixels during the imaging and digitization process. The value of *n* was determined empirically through plotting the discrepancy between the Fourier representation and target shape (residual sum of squares) with respect to increasing *n* and identifying where further increases in *n* does not result in meaningful improvements to the Fourier representation. This process was completed for 100 randomly selected cells (Supplemental Fig. 1). We determined that *n* = 15 (62 coefficients total) and *n* = 5 (22 coefficients total) were sufficient to faithfully reconstruct cellular bodies and nuclei, respectively.

### Quantification of αSMA stress fiber structures

After achieving mathematical descriptors of VIC cell body and nuclear shape, we developed complementary analytical methods to characterize αSMA stress fibers. Specifically, we computed coherence (*c*) which reflects the local degree of anisotropy^26–28^ and is a sensitive metric of αSMA stress fiber organization because of their fibrous nature. To compute *c*, we first computed the *x-* and *y-*gradients (*I*_*x*_ and *I*_*y*_, respectively) of αSMA fluorescent images on a pixel-by-pixel basis using the Sobel-Feldman operator^29^. Next, we assembled the structure tensor *S*

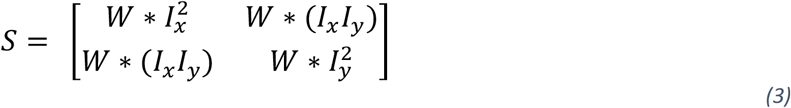

where *W* is a Gaussian smoothing kernel with a standard deviation of 4 and the symbol * denotes convolution. Eigendecomposition of S was performed to obtain Eigenvalues (*λ*_1_, *λ*_2_) and Eigenvectors (***e***_1_, ***e***_2_). If *λ*_1_=*λ*_2_, then the gradient has no preferred direction which can occur in the case of rotational symmetry or homogeneity (Fig. 2a, left-most panel). If *λ*_1_>*λ*_2_, then *λ*_1_ represents the greatest gradient (i.e., change in intensity) squared at a given pixel that is oriented along ***e***_1_ whereas *λ*_2_ represents the gradient squared along the orthogonal direction ***e***_2_ (Fig. 2a, middle three panels). In the case that *λ*_1_>0 and *λ*_2_=0, the structure in the image is completely aligned and is locally one-dimensional (Fig. 2a, right-most panel). The relative difference in magnitude between *λ*_1_ and *λ*_2_ reflects the local degree of anisotropy and is quantified by *c*

**Fig. 2:**
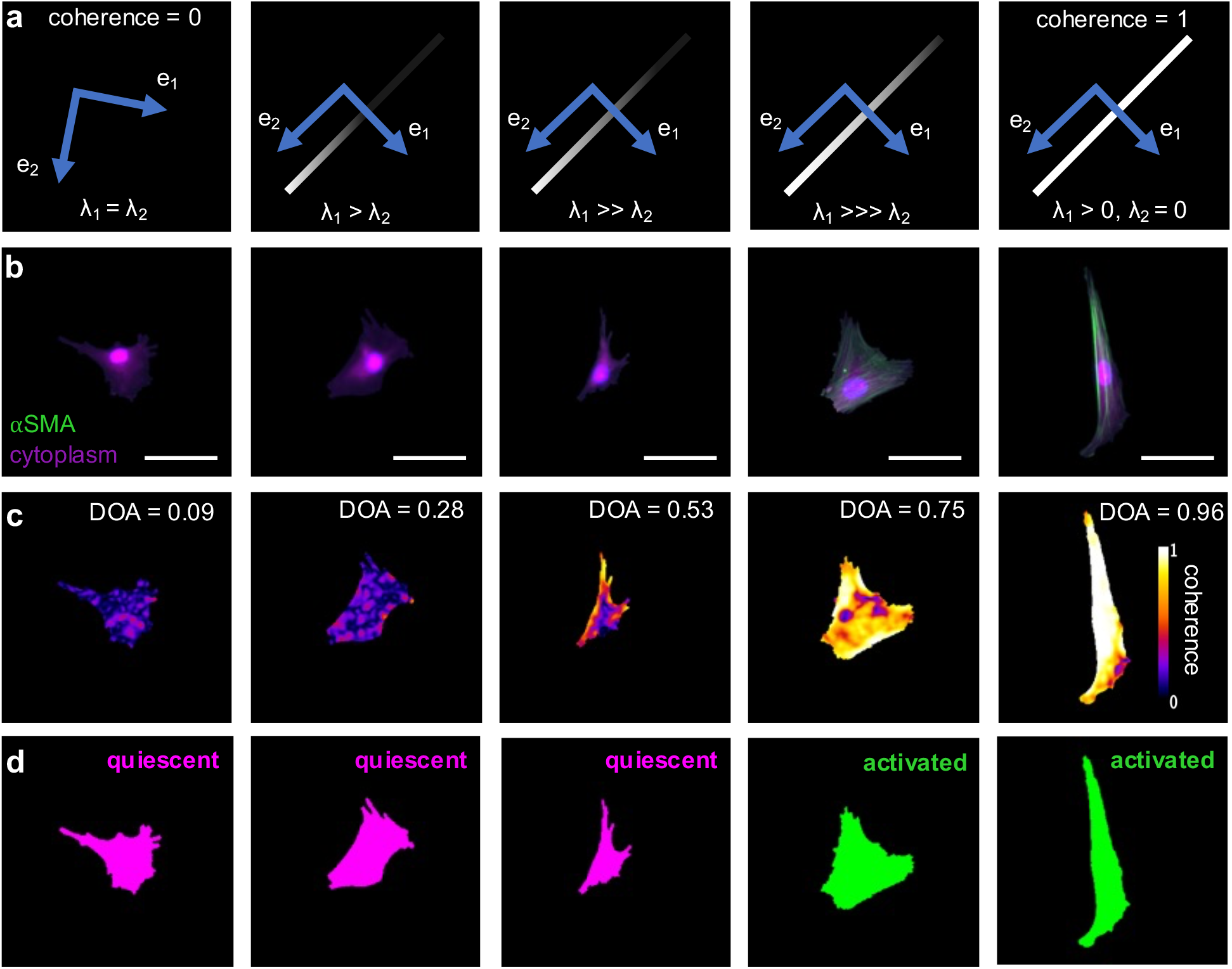
Quantification of αSMA stress fiber architecture. **a**, Schematic demonstrating the computation of coherence. Coherence, at a point, increases as the greatest intensity gradient (λ_1_) becomes greater than the orthogonal intensity gradient (λ_2_). For a fiber, λ_1_ is oriented along the cross-fiber direction (**e**_1_) whereas λ_2_ is oriented along the preferred fiber direction (**e**_2_). In the case of homogenous intensity values (left most panel), the directions for **e**_1_ and **e**_2_ are random but orthogonal. **b**, Representative immunostaining images depicting VICs with increasing levels of αSMA expression and anisotropy from left to right. **c**, Degree of anisotropy (DOA) is computed for the VICs shown in panel (**b**) by taking the median of the coherence values at pixel locations comprising the VIC body. **d**, Manual grading of the VICs shown in panel (**b**) as quiescent or activated. Stained images: green - αSMA, magenta – cytoplasm. Scale bars: 50 *µ*m.

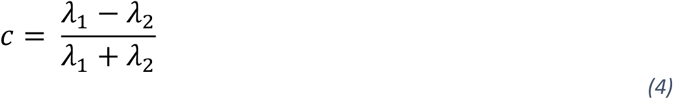

whose value ranges from 0 to 1. We demonstrate that *c* is resistant to changes in absolute fluorescence intensity, distance from the focal plane, and pixel resolution (Supplemental Fig. 2), highlighting its versatility in various imaging scenarios. Moreover, *c* is a continuous variable whose limits represent completely diffuse αSMA staining (*c* = 0) and discrete αSMA stress fibers (*c* = 1) (Fig. 2a). In this analysis, *c* provides a quantifiable measure of the amount of αSMA organized in stress fibers (Fig. 2b), thereby providing a continuum measure of the quiescent to activated VIC phenotypic spectrum. The computed values of *c* were large in regions with discrete αSMA stress fibers and low in regions with no αSMA stress fibers (Fig. 2b&c). Since *c* is computed on a per-pixel basis, we sought a single number to represent the overall degree of anisotropy (DOA) of a given cell. To this end, the median *c* value was computed for each VIC to represent the DOA. We then compared a currently prevalent manual grading system of activated and quiescent VICs with our calculated DOA. High values of DOA correspond with activated VICs, while quiescent VICs have low values of DOA (Fig. 2c&d), thus providing confidence that DOA can distinguish between traditionally graded VIC phenotypes while offering a refined and quantitative approach. Collectively, this quantitative method characterizes αSMA stress fiber expression while avoiding manual grading of VIC phenotypes into binary classifications.

### Extraction of morphological features that correlate with VIC activation

Prior methods to quantify and describe cellular morphologies have relied on domain knowledge to select and manipulate raw data into intuitive and meaningful features such as cell area, perimeter, solidity, circularity, aspect ratio, and curvature^20^. In the method presented, experimenter subjectivity is minimized during the process of morphological feature extraction and its subsequent correlation to αSMA anisotropy which reflects the activated phenotype. The methodology was first trained and validated on a dataset generated by seeding VICs on soft (E= ∼5 kPa) (Fig. 3a) and stiff (E= ∼13 kPa) (Fig. 3b) hydrogel matrices that mimic healthy and fibrotic valve tissue matrix and are known to induce quiescent and activated VIC phenotypes, respectively^30–32^. A single metric, the degree of anisotropy (DOA), was used to quantify the amount of αSMA in discrete stress fibers present within the VICs which reflects their level of myofibroblast activation (Fig. 2b&c). Next, feature vectors were generated for every cell, which were comprised of the Fourier coefficients describing the cell body and nuclei and the associated DOA value (Supplemental Table 1). The feature vectors were standardized and concatenated into a single array, and then principal component analysis (PCA) was performed. PCA showed principal component (PC) axes in which the VICs are most variant. Visualization of the Eigenshapes, which show the greatest shape variation along orthogonal PC axes, revealed morphological features that co-varied with DOA (Fig. 3c). This process allowed for physical interpretation of morphological features that are most indicative of the activated phenotype. For example, we interpreted the morphological features of PCs 1, 2, and 3 as area, length, and aspect ratio, respectively (Fig. 3d). We computed the probability density estimates (PDE) for VICs seeded on soft and stiff hydrogels for every PC and found that on stiff, activation-promoting hydrogels, VICs showed larger PC scores along PCs 1 and 3 compared to VICs seeded on soft, quiescent-promoting hydrogels (Fig. 3e). No appreciable difference was observed between the soft and stiff condition for PC 2. To assess the practical significance of the observed differences, the effect size for each PC axis was represented using Cohen’s d^33,34^. A Cohen’s d value of 0.94 was observed between the soft and stiff condition along PC1 (large to very large effect size) and 0.61 along PC3 (medium to large effect size, Supplemental Table 2). All other PC axes showed a Cohen’s d < 0.5 (lower than medium effect size) and thus were not considered for further analysis. Taken together, these results suggest cell and nuclear area and aspect ratio are morphological features that are correlated with VIC activation whereas cell and nuclear length are not (Fig. 3e). Furthermore, there was no significant correlation observed between cell area and aspect ratio (*r* = 0.16) nor between nuclear area and aspect ratio (*r* = −0.10), suggesting that they are orthogonal morphological features that should be considered separately. We observed experimentally imaged cellular morphologies (Fig. 3f) that aligned closely with the Eigenshapes generated for PCs 1-3 (Fig. 3c). This finding validates that the current approach produces realistic Eigenshapes and highlights its relevance toward experimentally observed cellular geometries. In addition, the identified cells underline the innate heterogeneity of VICs that can be observed *in vitro*.

**Fig. 3:**
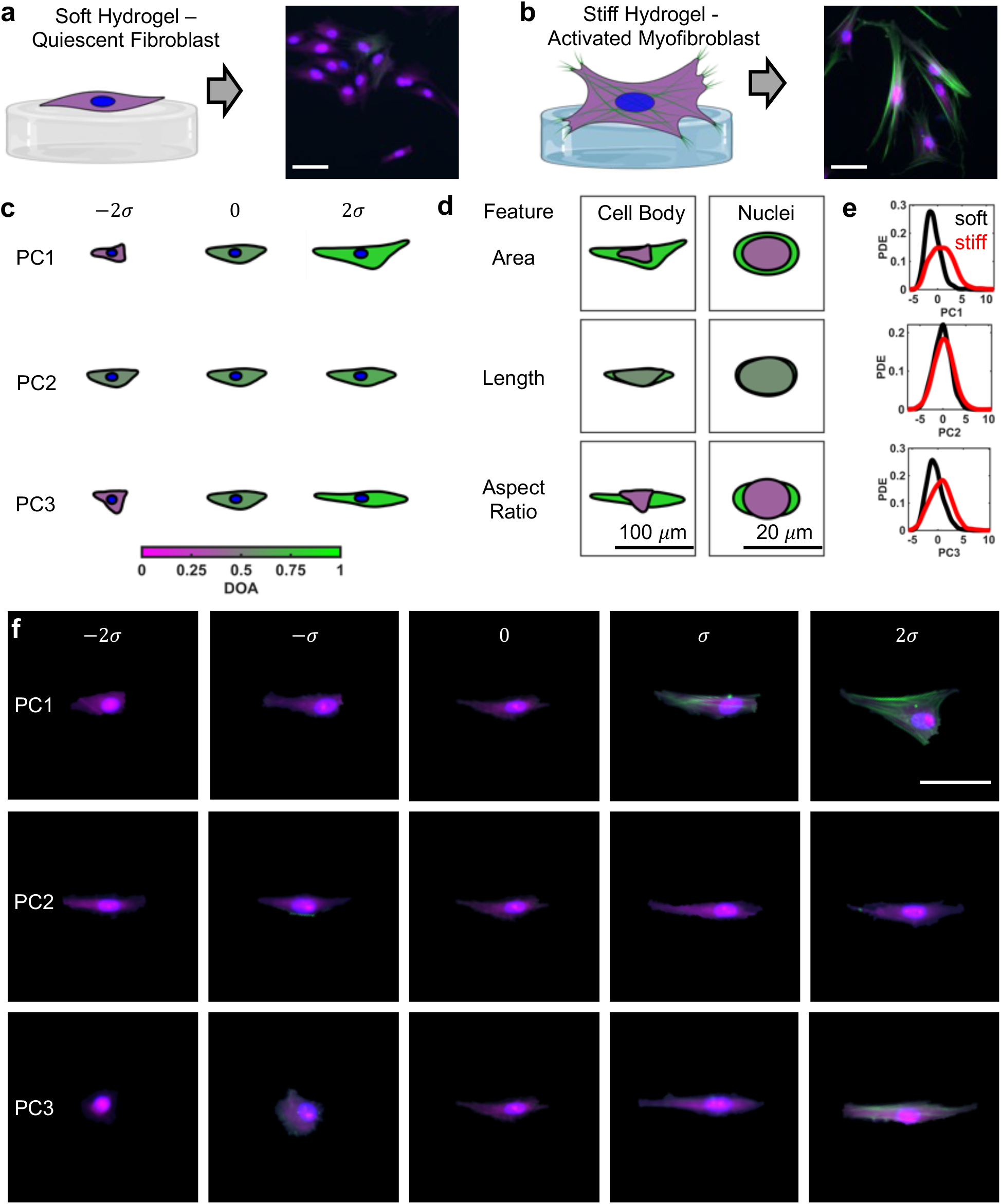
An approach combining morphological and αSMA expression data to reveal relevant morphological features to demarcate VICs seeded on engineered hydrogel materials with tunable mechanical properties. **a**, VICs seeded on top of mechanically soft hydrogels exhibit a quiescent phenotype characterized by limited expression of αSMA stress fibers. **b**, VICs seeded on top of mechanically stiff hydrogels transition to an activated myofibroblast phenotype characterized by high expression levels of αSMA. **c**, VIC body and nuclear Eigenshapes computed along principal components (PCs) 1-3 and colored with the corresponding DOA values. **d**, VIC body and nuclear Eigenshapes at the extremes (−2*σ* and 2*σ*) are superimposed and colored with the corresponding DOA values for PCs 1-3 accompanied with a physical interpretation of the primary morphological feature that varies along each PC. **e**, Probability density estimates (PDEs) depicting the PC score distributions for VICs seeded on soft (black) and stiff (red) hydrogels for PCs 1-3. **f**, Representative segmented VICs that closely align with Eigenshapes along principal components 1-3 in panel (**c**). Stained images: green - αSMA, magenta – cytoplasm, blue-nuclei. Scale bars: 50 *µ*m. Schematics created with BioRender.com.

### Selection and validation of features that can grade VIC phenotypes

PCA and generation of Eigenshapes along the PC axes (Fig. 3) identified that cell and nuclear area and aspect ratio covaried with DOA and were potential morphological features that could be used to grade quiescent and activated VICs (Fig. 4a). For each identified feature and DOA variable, logistic regression models were developed to grade VIC phenotypes based on morphology and αSMA anisotropy, respectively (Fig. 4b). In addition, we trained a multivariable logistic regression (MLR) model that combined the effects of each identified feature and DOA. For visualization of the MLR, we varied the parameters cell area and DOA, which differed greatly between quiescent and activated VICs (Supplemental Table 3), while keeping the remainder of the features constant and equal to the mean value observed for the VICs analyzed in the present study. Logistic regression models were employed due to their simplicity and interpretability. Specifically, the coefficients of the trained models inform on the appropriate threshold value to grade VICs for each identified feature (Supplemental Table 4). Moreover, the coefficients of the MLR model inform on the relative contribution or importance of each feature in a combined approach to grade VICs (Supplemental Table 5). Grading was accomplished by setting a threshold value (red star in Fig. 4b, Supplemental Table 4) for each feature that was defined by an activation probably ≥50%. To assess the performance of individual binary classifiers at varying threshold values, receiver operating characteristic (ROC) curves were generated for the features and the MLR model (Fig. 4c). ROC curves allowed for visual inspection of the area underneath the curve (AUC) which serves as a single quantitative descriptor of the predictive capability of a given variable. The AUC for cell area (0.73), nuclear area (0.76), DOA (0.90), and the MLR model (0.92) were high, thus identifying these as suitable variables for grading VIC phenotypes. In contrast, the AUC for cell (0.60) and nuclear (0.60) aspect ratio were relatively lower (Supplemental Table 6) and thus less predictive of VIC phenotypes. Of further note, activated VICs had significantly larger cell and nuclear areas and higher values of DOA (Fig. 4d), which all showed Cohen’s d values greater than 0.8 (large to very large effect sizes, Supplemental Table 3). The largest effect size was observed in the MLR model (Cohen’s d = 2.22, huge effect size). No appreciable differences were observed between activated and quiescent VICs in terms of cell and nuclear aspect ratios (Fig. 4d) and their effect sizes were considered small to medium with a Cohen’s d less than 0.5 (Supplemental Table 3). Overall, the training of logistic regression models allows for the grading of VICs as quiescent or activated based on morphological characteristics, αSMA anisotropy, or both. Extended analysis with ROC curves assesses the overall performance of each model and showed that some features (cell area, nuclear area, DOA, and the MLR) are better predictors of VIC phenotype than others (cell and nuclear aspect ratio). However, all features performed better than a random classifier which is denoted by the gray-dotted lines in Fig. 4c.

**Fig. 4:**
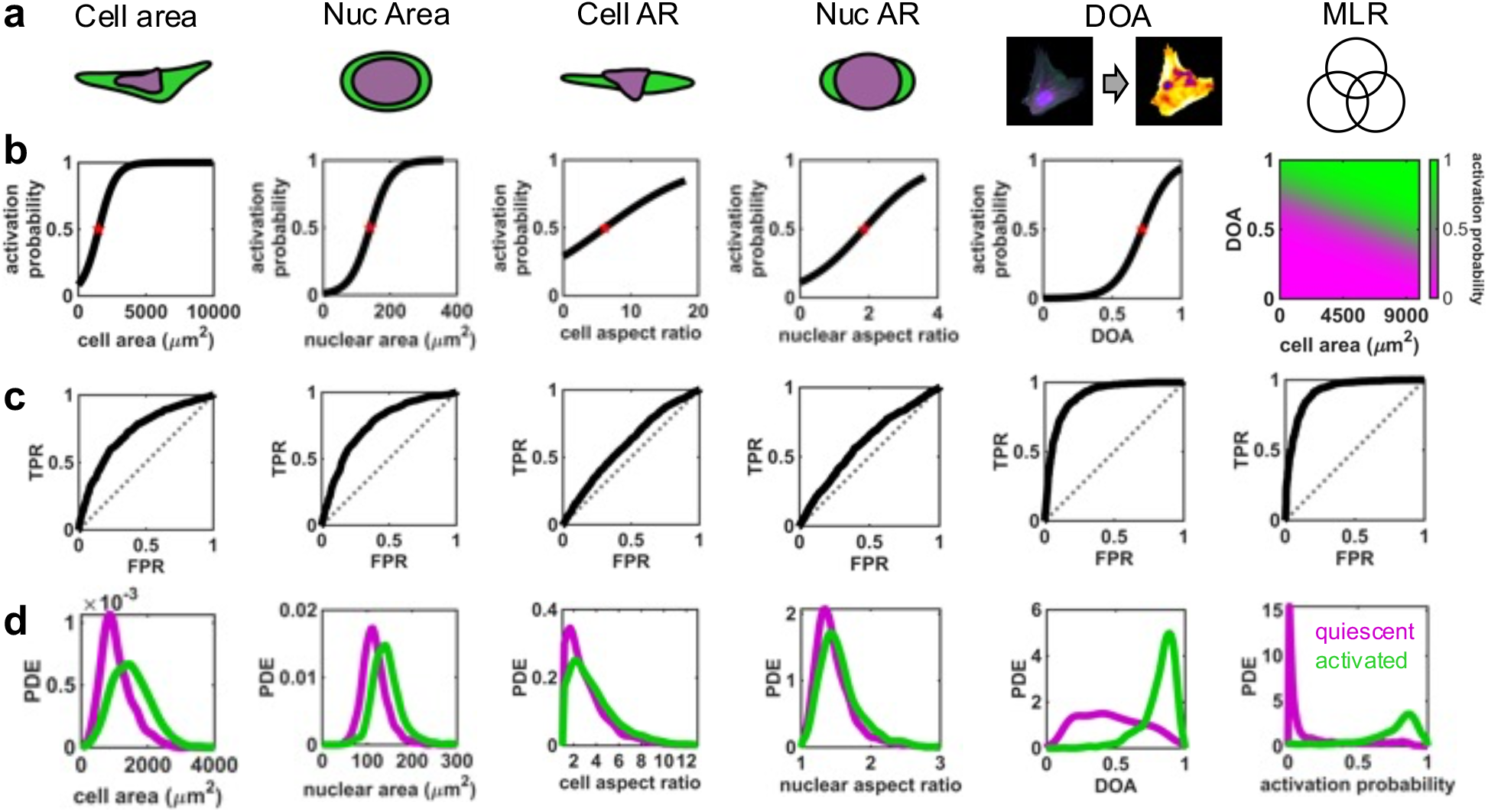
Efficacy of selected features toward demarcating quiescent and activated VICs induced by seeding on soft and stiff hydrogels. **a**, Visualization of selected features that most likely vary between quiescent and activated VICs. The unity symbol denotes a multivariable logistic regression (MLR) model which considers all selected features. **b**, Logistic regression curves depicting the activation probability of VICs versus the selected features. The red stars denote the quantitative threshold value where the activation probability of VICs ≥ 0.5. **c**, Receiver operating characteristic curves generated by plotting the true positive rate (TPR) against the false positive rate (FPR) at ever-increasing threshold values of the selected features. The ROC curve for a random classifier is shown by the gray-dotted line. **d**, Probability density estimates (PDEs) depicting the distributions of the selected features for quiescent (magenta) and activated (green) VICs. From left to right, the columns represent results for cell area, nuclear area, cell aspect ratio, nuclear aspect ratio, DOA, and the MLR model.

To validate the performance of the logistic regression models generated for each feature (Fig. 4b), we compared their predictions with manual phenotypic grading of fluorescent imaging datasets comprised of VICs stained for their cytoplasm, nuclei (DAPI), and αSMA stress fibers (Fig. 5a). A k-fold cross validation (k = 10) was used, in which the data was split into 10 random groups with 9 groups used to train the logistic regression models and the 10th serving as the validation data set (Fig. 5b). This process was iterated 10 times so that each group served as the validation data set exactly once. The percentage of cell phenotypes that were correctly predicted in the validation data set by the logistic regression models was recorded for every iteration (Fig. 5c). The validation showed that cell and nuclear area are good predictors of VIC phenotype with an accuracy of 69±3% and 70±3%, respectively; however, the cell and nuclear aspect ratios were less predictive with an accuracy of 60±3% and 61±3%, respectively. The DOA variable had the highest predictive capability among the single variables tested with an accuracy of 83±2%.

**Fig. 5:**
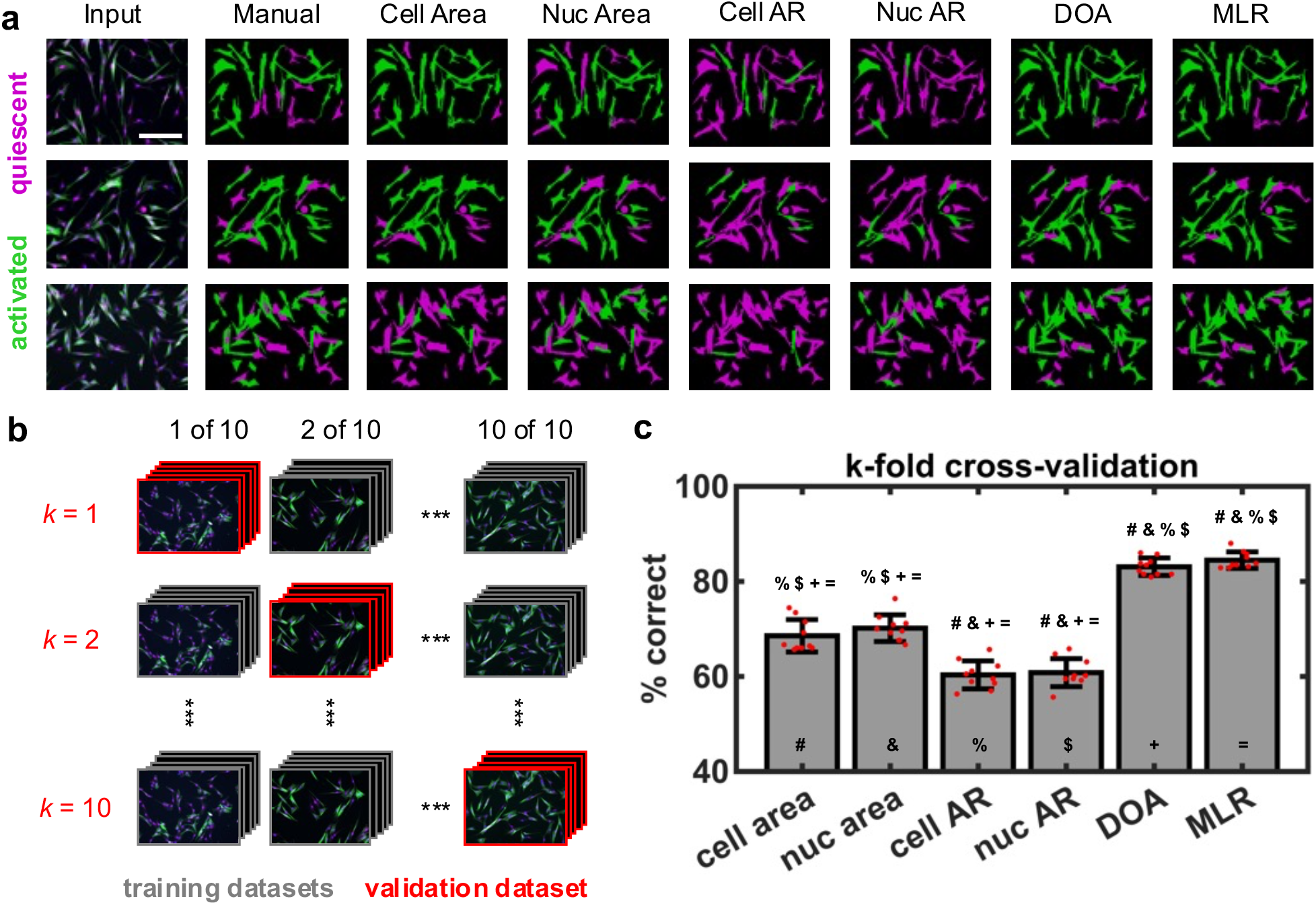
Validation of selected features toward grading activated and quiescent VICs. **a**, Three representative examples of the grading process depicting the input image, manual grading, grading done by quantitative thresholding of selected features (cell area, nuclear area, cell aspect ratio (cell AR), nuclear aspect ratio (nuc AR), and DOA) and grading done via a multivariable logistic regression (MLR) model that combines information from all the selected features. Quantitative thresholds were determined via logistic regression. **b**, To assess the predictive capabilities of the selected features, *k*-fold cross-validation was performed in which the dataset was randomly partitioned into *k* equal sized subsets. Of the *k* subsets, *k-1* subsets were used to train logistic regression models and the remaining subset was used as the validation subset. The cross-validation process was repeated *k* times, with each of the *k* subsets serving as the validation subset exactly one time. For this analysis, *k* = 10. **c**, Bar plots showing compiled *k*-fold cross-validation results depicting the average percentage of VICs graded correctly for each selected feature and the MLR when compared to manual grading. The error bars denote standard deviation. Each feature is represented with a symbol. The symbols on top of each boxplot denote the features that are significantly different from the current feature as determined by a one-way ANOVA (p < 0.01). Stained images: green - αSMA, magenta – cytoplasm, blue-nuclei. Scale bars: 200 *µ*m.

Beyond assessing the predictive capabilities of individual features, we also assessed the combination of all features on predicting VIC phenotypes using a MLR model (Supplemental Table 5). Overall, the MLR model showed the best predictive capabilities with an accuracy of 84±2%; however, combinations of features did not result in any significant improvements in the predictions compared to the DOA variable alone.

### Performance of models on test dataset

The ability of the logistic regression models, which were trained on VICs seeded on soft and stiff hydrogels, to grade VICs from a test dataset was assessed using a separate cohort of VICs seeded on top of stiff, activation-promoting hydrogels treated with the small molecule drug 5-azacytidine (AZA)^35^ (Fig. 6a). AZA has been reported to be protective against fibrosis by increasing the expression of phosphatase and tension homolog^36–40^. The logistic regression models for cell area, nuclear area, DOA, and the MLR model showed excellent performance in predicting the phenotype of VICs in the test dataset with accuracies of 75%, 72%, 74%, and 84%, respectively (Fig. 6b&c). The logistic regression models for cell and nuclear aspect ratio showed reduced performance with an accuracy of 61% and 55% in the test dataset, respectively. Based on manual grading, 88% of VICs seeded on soft hydrogels were quiescent while 12% were activated, 32% of VICs seeded on stiff hydrogels were quiescent while 68% were activated, and 73% of VICs seeded on stiff hydrogels with AZA treatment were quiescent while 27% were activated (Fig. 6d). Of note, the predictions made by the MLR on the soft and stiff dataset as well as on the test dataset (soft - 92% quiescent, 8% activated; stiff - 23% quiescent, 77% activated; AZA - 72% quiescent, 28% activated) largely resembled manual grading, highlighting the ability of the current approach to recapitulate traditional methods while employing quantitative and data-driven methodologies. When taken together, these results highlight that VICs on stiff hydrogels treated with AZA closely resemble quiescent VICs on soft hydrogels, suggesting that AZA may protect against fibrosis by inhibiting myofibroblast activation. In addition to confirming the ability of the models to make predictions on a test dataset, these results showcase the utility of the present approach in drug-screening applications toward predicting the activation landscape of VICs in response to candidate anti-fibrotic drugs.

**Fig. 6:**
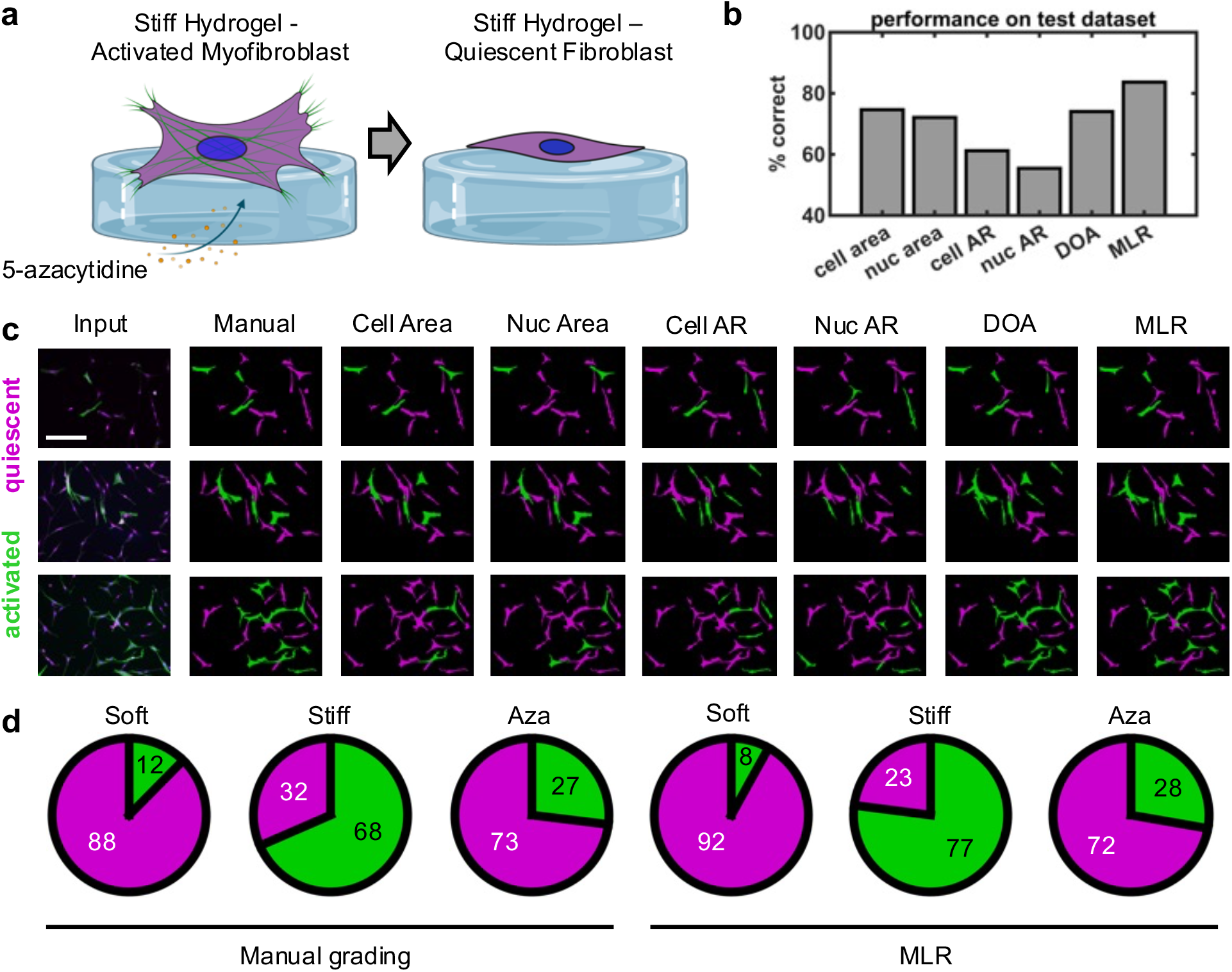
Model performance on test data set. **a**, VICs are seeded on top of stiff, activation-promoting hydrogel surfaces and treated with the small molecule drug 5-azacytidine, which inhibits myofibroblast activation. **b**, The percentage of VICs in the test dataset graded correctly for each selected feature and the MLR as compared to manual grading. **c**, Three representative examples of the grading process depicting the input image, manual grading, grading done by quantitative thresholding of cell area, nuclear area, cell aspect ratio, nuclear aspect ratio, DOA and the multivariable logistic regression (MLR) model. **d**, The percentage of quiescent or activated VICs when seeded on soft hydrogels, stiff hydrogels, and stiff hydrogels treated with 5-azacytidine (Aza) as determined by manual grading and the MLR model. Stained images: green - αSMA, magenta – cytoplasm, blue-nuclei. Scale bars: 200 *µ*m. Schematics created with BioRender.com.

## DISCUSSION

Traditional methods of grading myofibroblast activation have relied on manual and visual inspection for the presence of discrete αSMA stress fibers, with morphological characteristics being of secondary importance if not ignored altogether. Moreover, previous approaches have relied on subjective grading based on domain knowledge and qualitative assessment. Other studies that quantify fibroblast phenotypes commonly rely on metrices computed using absolute αSMA fluorescence intensity, which largely depends on sample preparation methods, hardware used (e.g., microscope), and imaging parameters. In the present study, we develop a method which can leverage and combine morphological and αSMA stress fiber information derived solely from fluorescent images to demarcate quiescent fibroblasts from activated myofibroblasts. The use of the DOA variable allows for a quantitative method to characterize discrete αSMA stress fiber expression, and therefore phenotypic activation, through measuring their anisotropy. The use of Fourier series to obtain coefficients to represent cell and nuclear morphological data in an abstract manner allows for data-driven approaches to discover morphological features that are correlated with myofibroblast activation. Of note, the current method is intended for use on images that are routinely taken to assess fibroblast phenotypes and requires no additional experimental effort.

DOA performed the best among the individual variables used to predict VIC phenotypes (Fig. 5c). This finding is expected as DOA reflects the presence of discrete αSMA stress fibers, which is the main characteristic that is used to manually grade fibroblast activation. However, the current method was able to reveal morphological features that are indicative of VIC phenotypic activation as evidenced by both cell and nuclear area proving comparable in accuracy to DOA when grading the test dataset (Fig. 6b). The MLR resulted in the best predictions on the test data (Fig. 6), underscoring the utility of a quantitative grading scheme that combines both morphological and αSMA stress fiber information.

Traditionally, grading of fibroblast phenotypes is largely binary with quiescent and activated being the two main categories. In the context of the current study, individual features and a combined approach are used to perform binary grading to compare their predictive capabilities against the current gold standard: manual binary grading. However, fibroblast phenotypes likely reside on a spectrum. In appreciation of this fact, the developed methodology can also predict the activation probability of fibroblasts on a continuous scale (Supplemental Fig. 3). Looking forward, this capability could lead to the discovery of sub-phenotypes that lie between quiescence and phenotypic activation.

Fibroblast cells naturally reside within 3D microenvironments and the current methods can be extended to analyze 3D imaging data. DOA can be easily modified to detect out of plane αSMA stress fiber anisotropy while spherical harmonics (the 3D extension of Fourier series) can be used to describe 3D cellular shapes^25,41^. Moreover, the current methods are not limited to analyzing VICs and can be used to study other fibroblast cell types, as we have demonstrated with cardiac fibroblasts (Appendix A). In addition, the current methods can be extended to segment and analyze bright field images, presenting the opportunity to quantify fibroblast phenotypes in real time and longitudinally to assess dynamic phenotypic shifts. Lastly, the presented methodology has potential to augment and improve research in the field of fibrosis through the analysis of cellular and nuclear morphologies derived from histology of healthy and diseased patient samples. This analysis could include parameters which describe nuclear activity such as the chromatin condensation parameter^32,42^. In summary, our work establishes a method to learn how fibroblast morphologies change with respect to phenotypic activation and leverages this knowledge to quantitatively grade fibroblast phenotypes. Looking forward, the developed methodology can augment high-throughput screening of small molecule drug libraries by efficiently pinpointing drugs that induce a quiescent fibroblast phenotype and provide rational for further testing of candidate anti-fibrotic agents.

## ONLINE METHODS

### Seeding aortic valve interstitial cells on soft and stiff hydrogels

Primary porcine aortic valve interstitial cells (VICs) were isolated using previously published methods^43^. In brief, freshly isolated porcine aortic valve leaflets were obtained from Animal Biotech Industries and washed in Earle’s balanced salt solution (Sigma) supplemented with 1 *µ*g/ml amphotericin B (Gibco), 50 *µ*g/ml streptomycin (ThermoFisher), and 50 U/ml of penicillin (ThermoFisher). Next, the leaflets were digested within a collagenase solution comprised of 250 U/ml of type II collagenase (Worthington) dissolved within Earle’s balanced salt solution for 30 minutes. This primary digestion removed the valvular endothelial cells which were discarded. The leaflets were then transferred to a fresh collagenous solution and were incubated with gentle shaking for 60 minutes. Next, the solution containing the leaflets was vortexed for 5 minutes to help dislodge the VICs and was then poured through a 100 *µ*m cell strainer to collect the VICs. The isolated VICs were then pelleted and seeded within expansion media comprised of M199 media (ThermoFisher) supplemented with 10% Fetal Bovine Serum (FBS) (ThermoFisher), 1 *µ*g/ml amphotericin B, 50 *µ*g/ml streptomycin, and 50 U/ml of penicillin. The VICs were cultured until ∼80% confluency and detached using trypsin (ThermoFisher) for passaging, freezing, and experimentation.

VICs were seeded on soft (E = ∼2.8 kPa) and stiff (E = ∼13.4 kPa) hydrogels as previously described^44^ (Fig. 3a&b). In brief, poly(ethylene glycol) photodegradable diacrylate (PEGdiPDA; MW ∼3,400 Da) was co-polymerized with poly(ethylene glycol) monoacrylate (PEGA; MW ∼ 400 Da) and acrylamide diethylene glycol – diethylene glycol – glycine – arginine – glycine – aspartic acid – serine – glycine (Ac-OOGRGDSG) in PBS through a redox-initiated free radical polymerization. Hydrogel formulations were prepared with the following concentrations: 7.0 wt% PEGdiPDA, 6.8 wt% PEGA, 5mM Ac-OOGRGDSG, 0.2 M ammonium persulfate (APS), and 0.1 M tetramethylethylenediamine (TEMED). Approximately 100 *µ*m thick hydrogels were formed on acrylated glass coverslips after the addition of APS and TEMED. The hydrogels were allowed to cure for 6 minutes before being washed and stored in PBS prior to cell seeding. This formulation resulted in stiff hydrogels. To photodegrade and produce soft hydrogels, a separate cohort of stiff hydrogels were irradiated with 365 nm UV light at 10 mW/cm^2^ intensity for 6 minutes prior to cell seeding. VICs were seeded at a density of 15,000 cells/cm^2^ on 12 mm soft and stiff hydrogels within media comprised of M199 media supplemented with 1% FBS, penicillin-streptomycin, and amphotericin B for 72 hours before fixing and staining. To generate the test data set, a separate cohort of VICs seeded on stiff hydrogels was treated with 10 *µ*M of the small molecule drug 5-azacytidine for 72 hours before fixing and staining.

### Immunostaining and Imaging

Samples were fixed in 4% paraformaldehyde (PFA, Electron Microscopy Sciences) for 20 minutes at room temperature before washing twice with PBS for 5 minutes each. Next, the samples were permeabilized with 0.1% TritonX-100 (Fisher Scientific) in PBS for 20 minutes at room temperature. Then, non-specific binding was blocked for 1 hour using 5% bovine serum albumin (BSA, Sigma-Aldrich) dissolved in PBS before adding mouse anti-αSMA primary antibody (Cat. No. ab7817, Abcam) diluted in 5% BSA at a ratio of 1:200. The samples were then incubated overnight at 4°C. The next day, the primary antibody was removed, and the samples were washed twice with PBST (0.5% Tween-20 in PBS) and once in PBS. Then, a secondary antibody solution containing 1:200 goat-anti mouse Alexa 488 (Cat. No. A-11001, ThermoFisher), 1:1000 DAPI, and 1:5000 HCS Cell Mask Orange (Life Technologies) was added for 1 hour and then rinsed 2x with PBST and once with PBS before imaging. Imaging was performed using an Operetta high-content confocal microscope (PerkinElmer) with a 20x air objective resulting in 1360 x 1024 x 10 images at a resolution of 0.49 *µ*m/pixel in the x-y plane and 1 µm through the z-plane.

### Segmentation, centering, and alignment

The boundaries of cellular bodies were segmented using a marker-based watershed algorithm (Fig. 1a) that used segmented nuclei as markers (Fig. 1b). The segmentation was performed in the Harmony High-Content Imaging and Analysis Software (PerkinElmer). We briefly note that marker-based watershed segmentation can be achieved in a myriad of programs including Matlab, Python, and ImageJ and is not a point of novelty in the current study. Segmentation quality was manually checked and cells that were segmented poorly (e.g., erroneous cell boundaries resulting in incomplete segmentation or inclusion of neighboring cells) were discarded from further analysis. Moreover, dividing cells evidenced by nuclei splitting, were also discarded. In addition, only cellular bodies that were completely in the field of view (i.e., not touching the image boundary) were included in the analysis. The pixels comprising the segmented cell boundaries and the segmented nuclei boundaries were converted to cartesian points by multiplying the pixel location by the image resolution (0.49 *µ*m/pixel). To ensure that morphological analysis was invariant to cell location and pose, the cell bodies and nuclei were centered and aligned so that their centroids coincided with the origin and their longest axes were parallel to the *x*-axis (Fig. 1c). The longest axis was defined as the direction which accounted for the greatest variation in cell/nuclei boundary points as determined by conducting principal component analysis (PCA) on the cartesian points of the cell/nuclei boundary and then aligning the cell/nuclei along the first principal component axis. “Flipping” of the shapes along the vertical and/or horizontal axis was also permitted to ensure that the greatest portion of the cell/nuclei area was in the first quadrant. This type of shape-preserving transformation was performed to ensure that all large cellular structures (such as protrusions) lie in approximately the same location across all cells to facilitate consistent analysis.

### Fourier series expansion

We employed Fourier-based methods to obtain mathematical descriptions of cellular bodies and nuclei to quantify their morphologies^25^. Two parametric functions x(θ) and y(θ) are employed to relate the arc length around a given cell/nuclei boundary (θ) to the corresponding x- and y-Cartesian coordinates. Here, the arc length of the shape is normalized such that the total length is 2*π*. Essentially, this results in a one-to-one mapping of the original shape to a unit circle, ensuring that every cartesian x,y point corresponds to exactly one arc length θ in the range [0, 2*π*] (equations 1 and 2). In equations 1 and 2, the variables a_0_, a_n_, b_n_, c_0_, c_n_, and d_n_ are Fourier coefficients determined by solving overdetermined systems of linear equations. Equations 1 and 2 can be represented in matrix form as

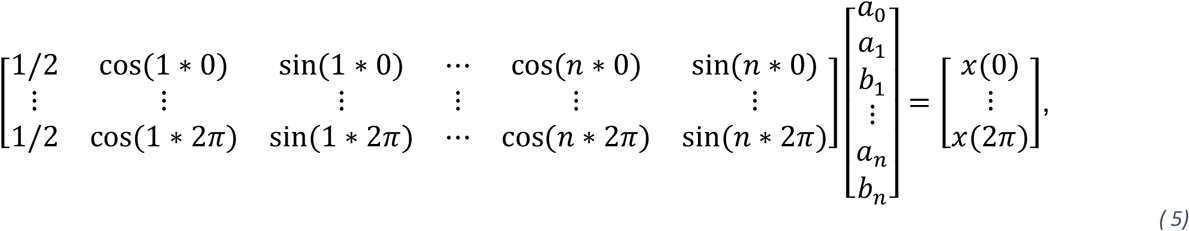

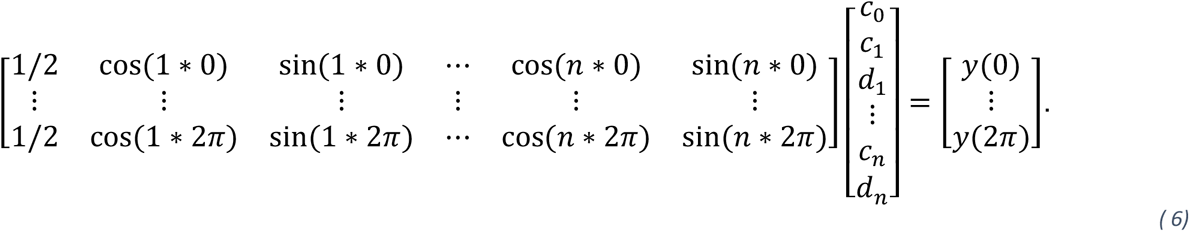

Here, *n* is the number of user-determined Fourier harmonics used to reproduce the shape. As *n* increases, the Fourier description converges to the target shape (Fig. 1d). Representing the boundaries of cell bodies and nuclei using Fourier series resulted in Fourier coefficients that contained information on cell and nuclei morphology (e.g., both size and shape) that can be used to analyze cellular bodies and nuclei in a data-driven manner.

### Computation of coherence and degree of anisotropy (DOA)

Coherence was computed to achieve a quantitative descriptor of αSMA stress fiber anisotropy or discreteness within imaged cells using previously described methods^26^ (Fig. 2a). We first computed the x- and y-gradients (*I*_*x*_ and *I*_*y*_, respectively) of αSMA images on a pixel-by-pixel basis using the Sobel-Feldman operator^29^ which consists of two 3x3 kernels

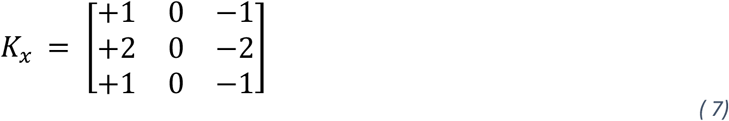

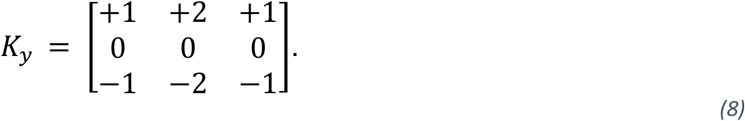

The convolution of *K*_*x*_ and *K*_*y*_ with an image *I* results in the gradients

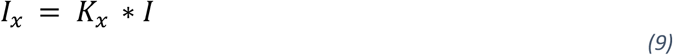

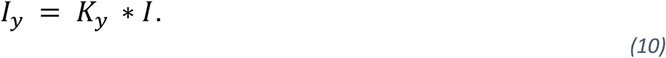

Here, the symbol * denotes the 2D convolution operation. Next, we assembled the structure tensor (equation 3). In equation 3, *W* is a Gaussian smoothing kernel with a standard deviation of 4 pixels and here again the symbol * denotes convolution. A standard deviation of 4 was chosen as the diameter of αSMA stress fibers in our fluorescence images spanned approximately 4 pixels. It is important to note that the standard deviation value will vary depending upon imaging parameters and the size (in pixels) of the biological structure of interest. Eigendecomposition of S was performed to obtain Eigenvalues (*λ*_1_, *λ*_2_) and Eigenvectors (***e***_1_, ***e***_2_). If *λ*_1_=*λ*_2_, then the gradient has no preferred direction which can occur in the case of rotational symmetry or homogeneity (Fig. 2a, left-most panel). If *λ*_1_>*λ*_2_, then *λ*_1_ represents the greatest gradient (i.e., change in intensity) squared at a given pixel which is oriented along ***e***_1_ whereas *λ*_2_ represents the gradient squared along the orthogonal direction ***e***_2_ (Fig. 2a, middle three panels). In the case that *λ*_1_>0 and *λ*_2_=0, the structure in the image is completely aligned and is locally one-dimensional (Fig. 2a, right-most panel). The relative difference in magnitude between *λ*_1_ and *λ*_2_ reflects the local degree of anisotropy. This is quantified by coherence (c), as shown in equation 4, whose value ranges from 0 to 1. The median coherence value of the pixels comprising the cell body was computed to achieve a single quantitative metric of the degree of anisotropy (DOA) of αSMA stress fibers (Fig. 2c). Traditionally, manual inspection of the presence of discrete αSMA stress fibers governed the grading of quiescent and activated fibroblasts (Fig. 2d).

### Concatenation and standardization of variables, principal component analysis, and generation of Eigenshapes

Feature vectors were established for each cell by combining the Fourier coefficients for the cell body and nucleus with the corresponding DOA value (Supplemental Table 1). The feature vectors were concatenated into an array such that every row contained information for one cell and every column represented a unique variable (either a Fourier coefficient or DOA). The variables were standardized as follows

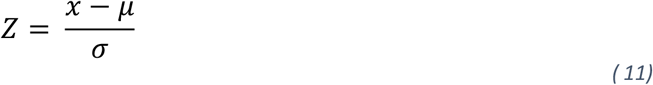

where *Z* represents the standard score, *x* is the observed value of the variable, *µ* is the mean of the variable, and *σ* is the standard deviation of the variable. Standardization of variables was performed to ensure that subsequent PCA was not affected by the magnitude of the variables. PCA was then performed to investigate the greatest sources of variation among the feature vectors which resulted in Eigenvalues and Eigenvectors (*U*). Next, principal component (PC) scores were computed using the following

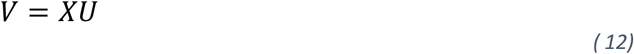

where *V* contains the resulting PC scores and *X* is the original data matrix containing the standardized feature vectors for every cell. Eigenshapes were generated along the principal axes by first taking the average of the PC scores across all cells analyzed. Then, the average PC score for the principal axis of interest was altered systematically from - 2*STD to +2*STD to produce Eigenshapes using

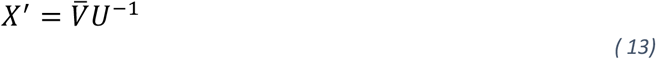

where 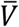 is the altered average PC scores and *X*^′^ is the recovered standardized feature vector. To generate the Eigenshape, *X*^′^ is first converted from Z-scores back to non-standardized variables to recover Fourier coefficients and DOA. The Fourier coefficients are used to generate the x-and y-cartesian coordinates of the Eigenshape using equations 1 and 2, respectively (Fig. 3c). The Eigenshape is colored based on its DOA value. Superimposing the -2*STD and +2*STD Eigenshapes along PC axes 1-3 allowed for physical interpretation of each PC axis and for the identification of features (Fig. 3d).

### Logistic regression models of features

Logistic regression models were trained for each feature (Fig. 4a) using a manually graded VIC data set (ground truth dataset) in which VICs were demarcated as quiescent (activation probability = 0) or activated (activation probability = 1). The training process resulted in models that predicted VIC phenotype based on individual features (Fig. 4b). The training process involved fitting the ground truth data set to the logistic function

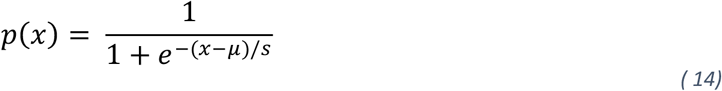

where *p*(*x*) is the probability of VICs being activated, *x* is a value of the feature of interest, *µ* is a location parameter such that *p*(*µ*) = 0.5, and *s* is a scaling parameter. In practice, the ground truth data set had activation probabilities of 0 or 1 whereas the fitted logistic regression model can produce continuous activation probabilities from 0 to 1. The coefficients for the logistic regression models were determined by minimizing the overall negative log-likelihood. To combine the effects of multiple features (e.g., cell area, DOA, cell aspect ratio) into a single model for predicting VIC activation, we trained a multi-variable logistic regression model

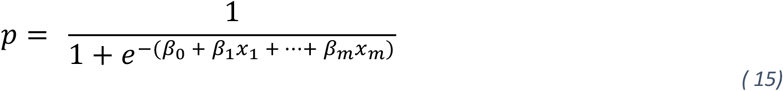

where *p* is activation probability, *β*_0−*m*_ are fitted parameters, and *x*_1−*m*_ are values of features. The coefficients were determined in similar fashion to those in equation 14 by minimizing the overall negative log-likelihood.

### Receiver operating characteristic curves

Receiver operating characteristic (ROC) curves^45^ were generated to visualize the predictive capability of each feature and the MLR model (Fig. 4c). This was accomplished by setting an ever-increasing threshold value for a specific feature of interest and computing the true positive rate (TPR) and false positive rate (FPR) at every threshold value. For the MLR, the probability at which a fibroblast is deemed activated was varied. The TPR and FPR were computed by comparing the grading of VIC phenotype by the current threshold value to the ground truth data set generated through manual grading. Next, the TPR was plotted against the FPR at every threshold value to achieve a visual representation of the feature’s predictive capability. In general, variables with high predictive capabilities will produce an ROC curve that approaches the top left corner of the plot whereas worse predictors will produce an ROC curve closer to the diagonal axis of the plot with the absolute worst predictors (i.e., random classifiers) forming a line of identity. The area under the curve (AUC) is computed to serve as a single quantitative descriptor of the predictive capability of each feature and the MLR model. AUC ranges from 0.5 (random classifier) to 1.0 (perfect classifier).

### k-fold cross validation

We performed k-fold cross validation to validate the ability of identified features to grade fibroblast phenotypes (Fig. 5b). Here, we chose k = 10. The entire data set was randomly separated into 10 equal groups where 9 of the groups were used to train the logistic regression models and the remaining group was used for validation. The percent of VICs in the validation data set that were predicted correctly as compared to manual grading was computed and served as an indicator of model accuracy (Fig. 5c). This process was iterated 10 times so that each group served as the validation group exactly once.

## Acknowledgements

We thank Dr. Tova L. Ceccato for the rat cardiac fibroblast imaging dataset. We acknowledge funding from the National Institutes of Health T32 HL007822 awarded to A.K., Helen Hay Whitney Foundation award number F1339 to K.B., American Heart Association 20PRE35200068 awarded to D.B. and National Institutes of Health R01 HL142935 and R01 HL132353 awarded to KSA.

## Author contributions

A.K., D.B., and K.S.A. designed and performed all the experiments. A.B. and A.K. performed image analysis, mathematical analysis, and data analysis. A.K., G.T., K.B., and K.S.A. contributed to writing and editing the paper.

## SUPPLEMENTARY INFORMATION

**Supplemental Fig. 1:**
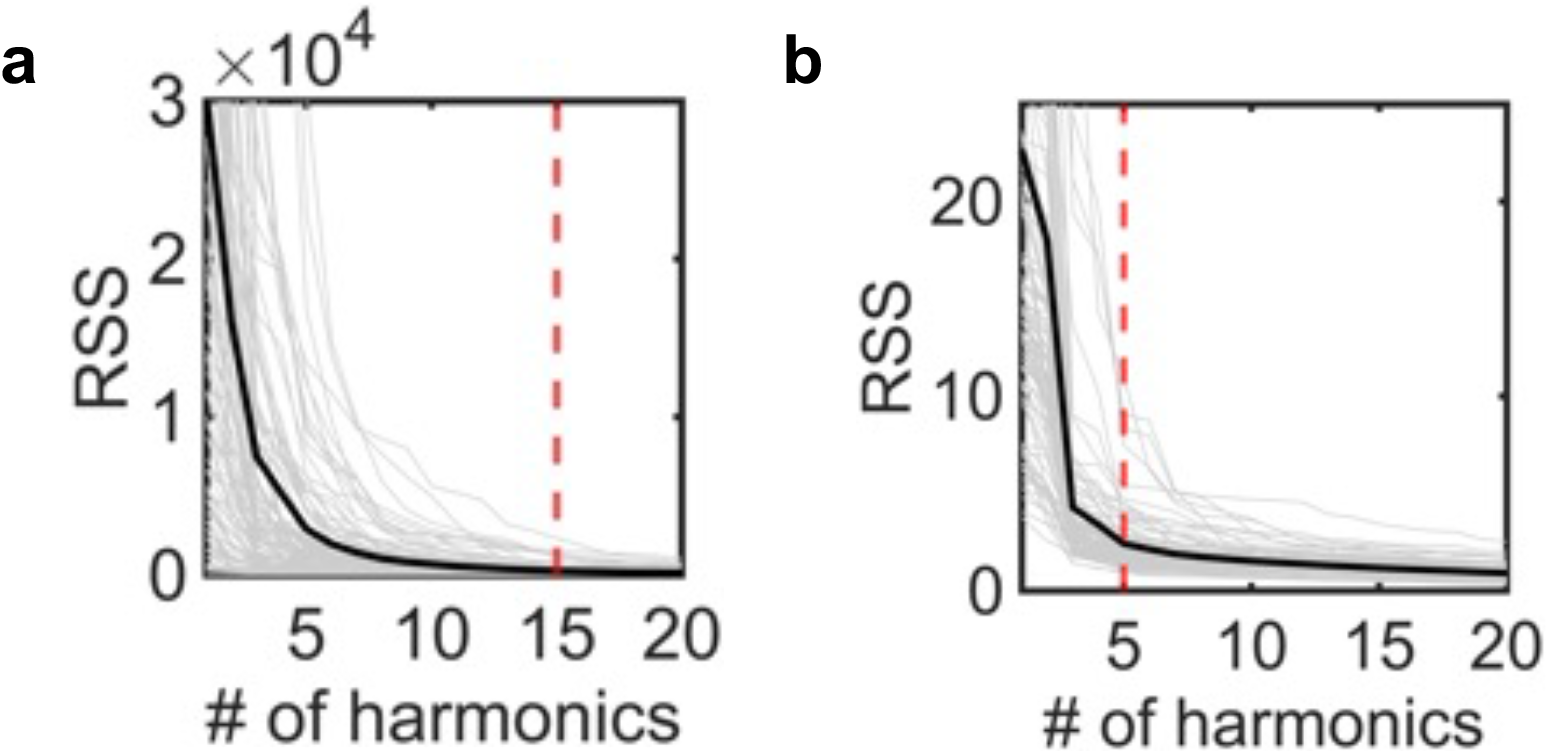
The appropriate number of Fourier series harmonics were chosen empirically to be **a**, 15 for cellular bodies and **b**, 5 for nuclear geometries based on the observation in which increasing the number of harmonics no longer produced an appreciable decrease in the residual sum of squares (RSS). The gray lines denote the results for individual cell bodies and nuclei. The black line denotes the mean of the gray lines. The red line denotes the empirical cutoff for the total number of harmonics used in Fourier representation of cell bodies and nuclei.

**Supplemental Fig. 2:**
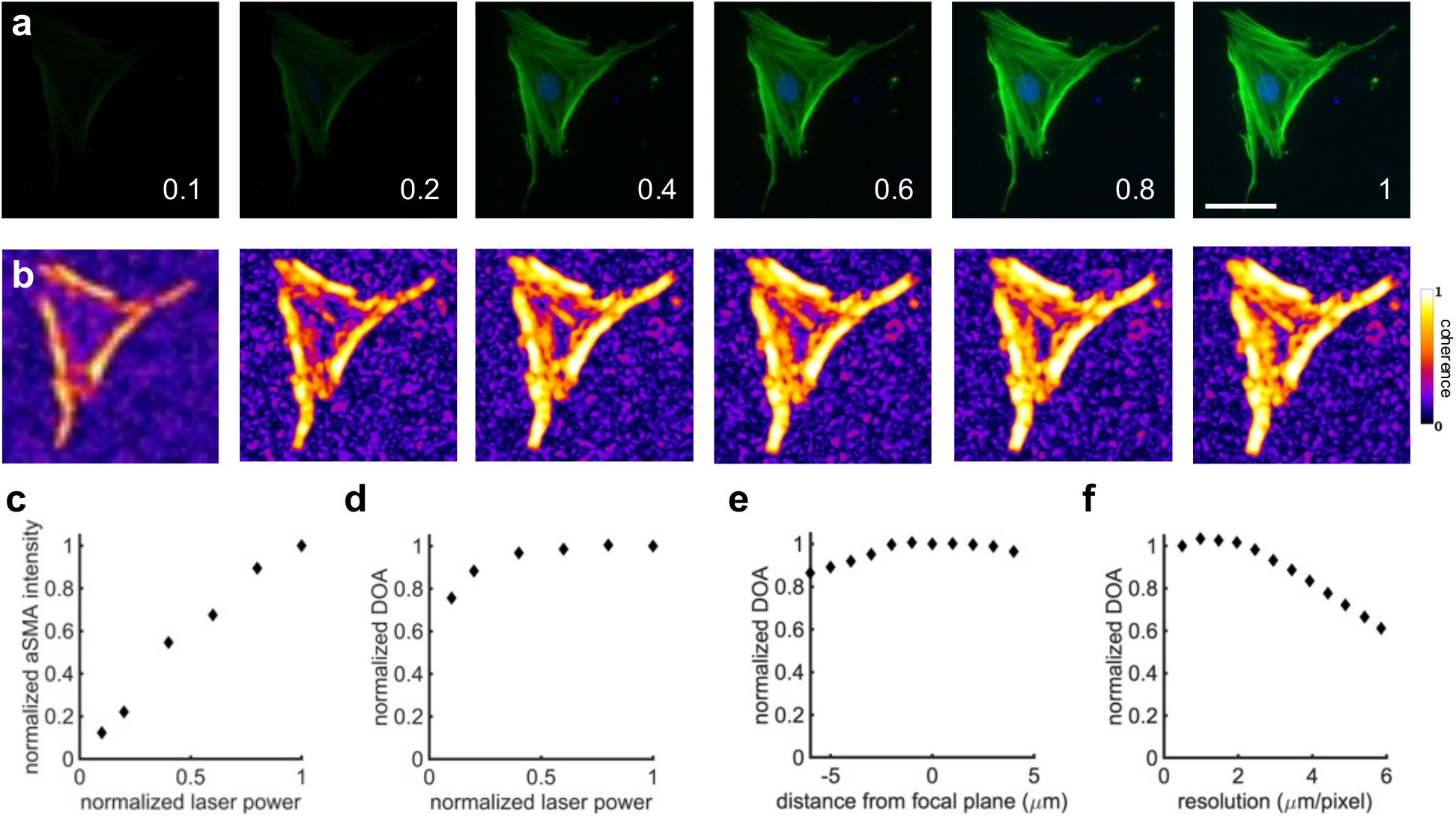
The effects of microscopy settings on DOA. **a**, Experimental images depicting a VIC imaged with increasing laser power. green - αSMA, blue – nuclei. **b**, Coherence computed for the corresponding images in panel (**a**), demonstrating that coherence is resistant to changes in laser power. **c**, The fluorescence intensity of αSMA is dependent upon laser power, highlighting a limitation of using fluorescence intensity dependent metrics to grade VIC activation. However, computed DOA values are resistant to changes in **d**, laser power and **e**, distance from the focal plane. **f**, DOA values are resistant to pixel resolution up to ∼2 µm/pixel. Stained images: green - αSMA, blue-nuclei. Scale bar: 50 *µ*m.

**Supplemental Figure 3:**
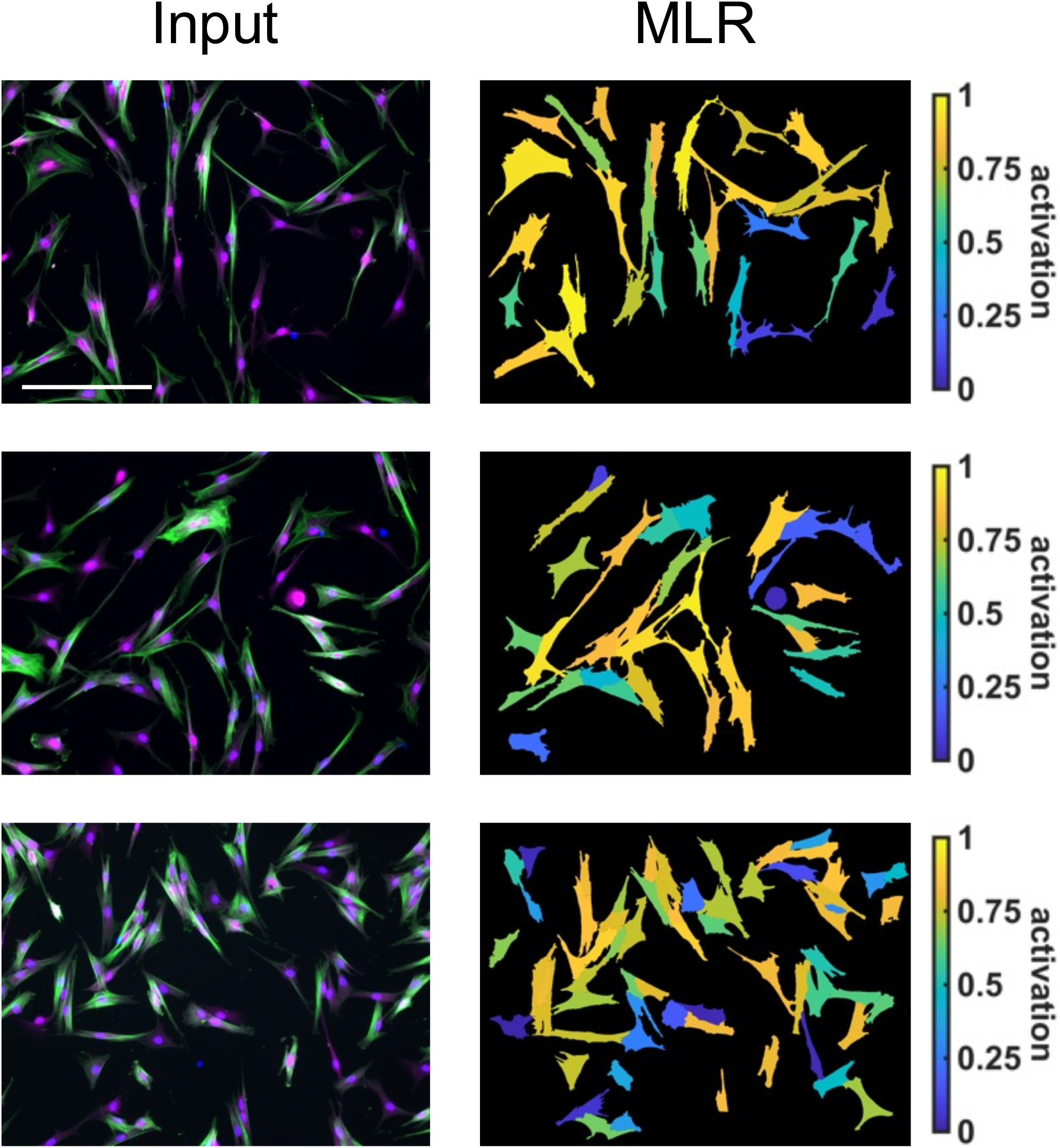
Three representative cases showing the ability of the multivariable logistic regression model to grade VIC activation on a spectrum. Stained images: green - αSMA, magenta – cytoplasm, blue-nuclei. Scale bars: 200 *µ*m.

**Supplemental Table 1:**
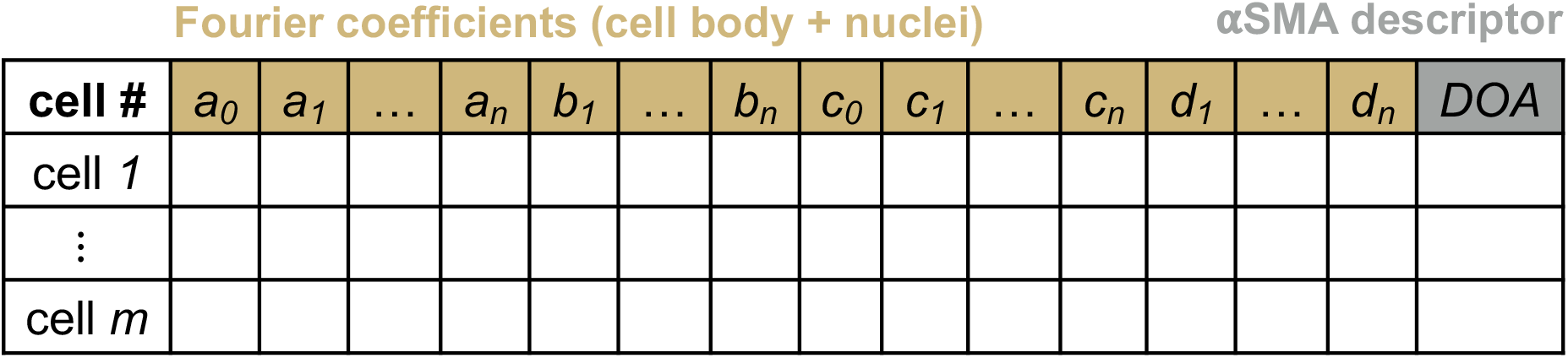
Demonstration of variable concatenation for principal component analysis. Both morphological (Fourier coefficients) and αSMA expression (DOA) information is combined. The columns denote a unique variable whereas the rows contain a unique vector per cell. All variables are standardized before PCA is performed.

**Supplemental Table 2:**
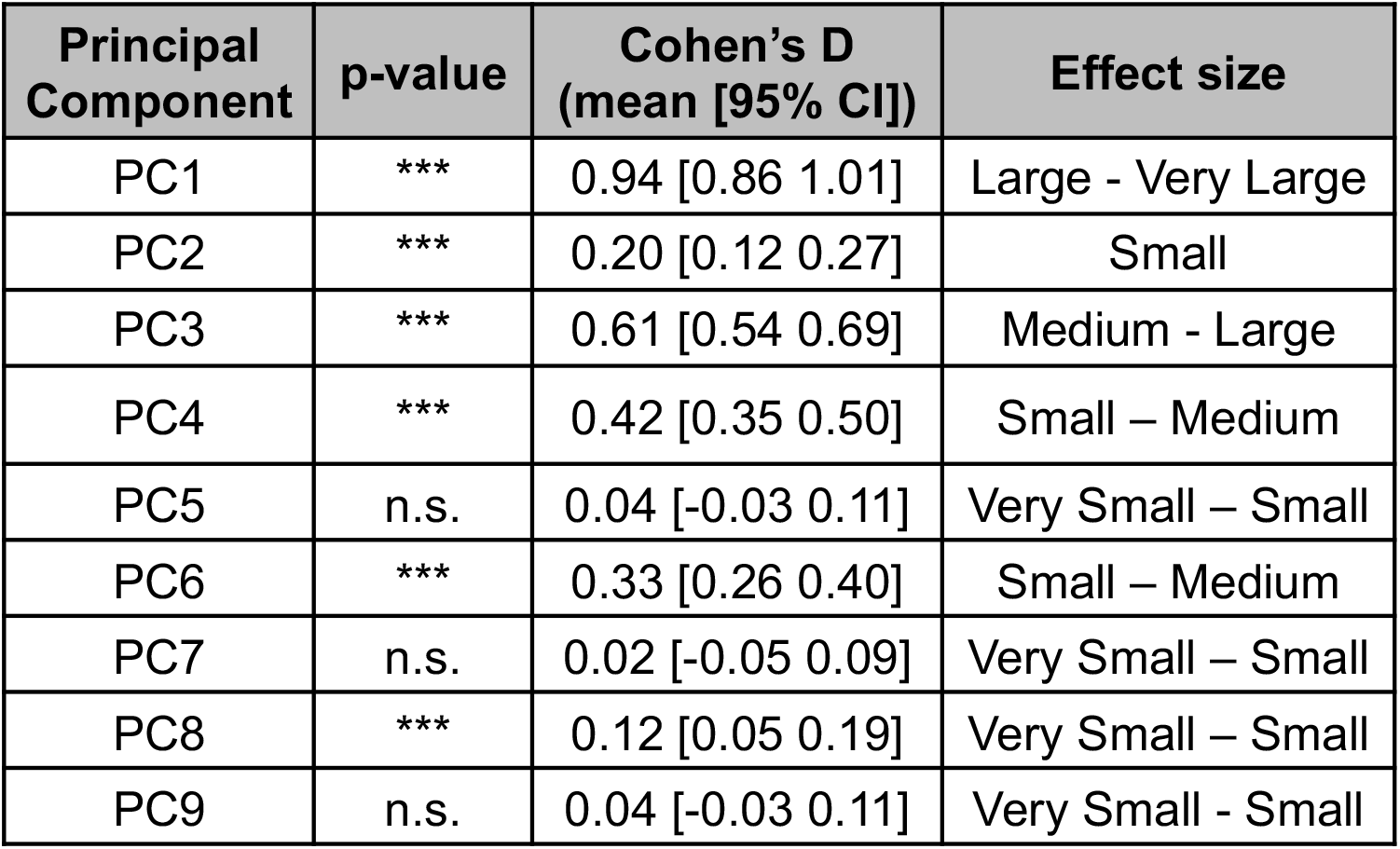
Statistical analysis of the principal component (PC) axes. P-values are presented for a two-sample t-test between PC scores of VICs seeded on soft and stiff hydrogels. Effect size is measured by Cohen’s D. n.s. – P > 0.05, *P ≤ 0.05, **P ≤ 0.01, ***P ≤ 0.001.

**Supplemental Table 3:** Statistical analysis of features. P-values are presented for a two-sample t-test between features of quiescent and activated VICs. Effect size is measured by Cohen’s D. n.s. – P > 0.05, *P ≤ 0.05, **P ≤ 0.01, ***P ≤ 0.001.

| Feature | p-value | Cohen's D<br>(mean [95% CI]) | Effect size |
| --- | --- | --- | --- |
| Cell area | *** | 0.82 [0.74 0.89] | Large – Very Large |
| Nuclear area | *** | 0.90 [0.83 0.98] | Large – Very Large |
| Cell aspect ratio | *** | 0.30 [0.23 0.38] | Small – Medium |
| Nuclear aspect ratio | *** | 0.33 [0.25 0.40] | Small – Medium |
| DOA | *** | 1.84 [1.76 1.93] | Very Large - Huge |
| MLR | *** | 2.22 [2.13 2.31] | Huge |

**Supplemental Table 4:**
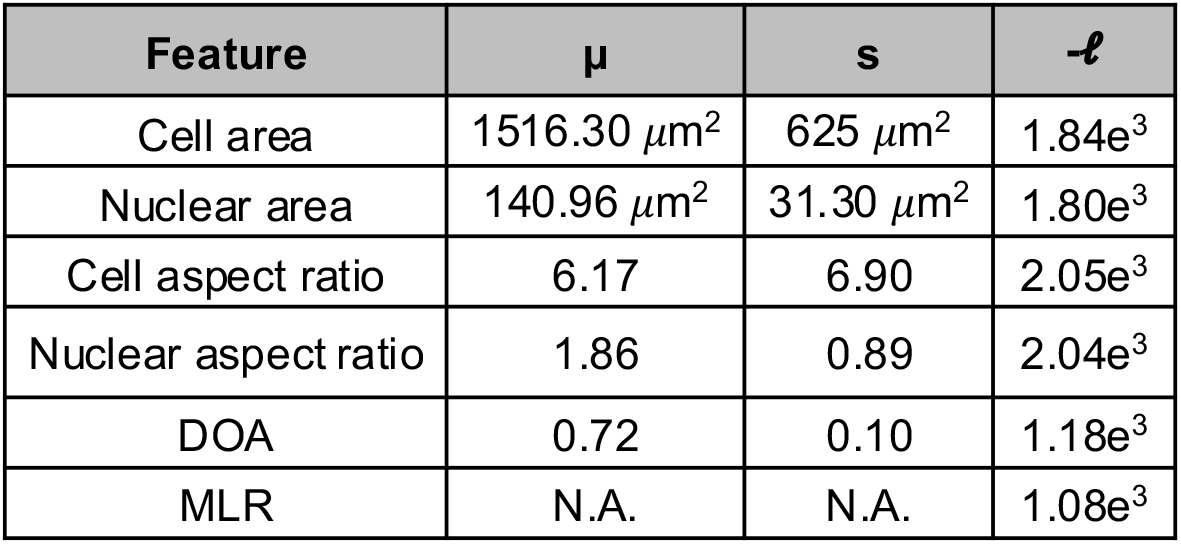
Compiled summary of logistic regression models showing the threshold value used to grade VIC phenotypes (*µ*), the scaling parameter (s), and the negative log-likelihood (-*𝓁*) for each model. A smaller value of -*𝓁* denotes a better fit.

| Feature | $\mu$ | s | $-\ell$ |
| --- | --- | --- | --- |
| Cell area | 1516.30 $\mu\text{m}^2$ | 625 $\mu\text{m}^2$ | 1.84e<sup>3</sup> |
| Nuclear area | 140.96 $\mu\text{m}^2$ | 31.30 $\mu\text{m}^2$ | 1.80e<sup>3</sup> |
| Cell aspect ratio | 6.17 | 6.90 | 2.05e<sup>3</sup> |
| Nuclear aspect ratio | 1.86 | 0.89 | 2.04e<sup>3</sup> |
| DOA | 0.72 | 0.10 | 1.18e<sup>3</sup> |
| MLR | N.A. | N.A. | 1.08e<sup>3</sup> |

**Supplemental Table 5:** Multivariable logistic regression model coefficients for the VIC dataset. n.s. – P > 0.05, *P ≤ 0.05, **P ≤ 0.01, ***P ≤ 0.001.

| Feature | Coefficient | Std. Error | P-value |
| --- | --- | --- | --- |
| Intercept ($\beta_0$) | -9.24 | 0.47 | *** |
| Cell area ($\beta_1$) | 4.04e-4 | 1.33e-4 | ** |
| Nuclear area ($\beta_2$) | 0.01 | 0.003 | *** |
| Cell aspect ratio ($\beta_3$) | -0.26 | 0.03 | *** |
| Nuclear aspect ratio ($\beta_4$) | 0.54 | 0.21 | * |
| DOA ($\beta_5$) | 10.21 | 0.42 | *** |

**Supplemental Table 6:** Area under the curve (AUC) for each feature and the MLR model. A higher AUC value denotes better model performance.

| Feature | AUC |
| --- | --- |
| Cell area | 0.73 |
| Nuclear area | 0.76 |
| Cell aspect ratio | 0.60 |
| Nuclear aspect ratio | 0.60 |
| DOA | 0.90 |
| MLR | 0.92 |

## Appendix A Extension of methodology to grade cardiac fibroblast phenotypes modulated through biochemical means

### General Overview

The ability of the present approach to be extended to grade other fibroblast cells was assessed using cardiac fibroblasts (CFs) treated with and without transforming growth factor beta (TGF-β), a compound widely known to elicit phenotypic activation^46,47^. First, we assessed the extent to which the VIC-trained models may generalize beyond their training data and how they perform on a different type of fibroblast. Then we compare the performance of the VIC-trained models against models that are directly trained on the CF dataset.

#### Culturing CFs with and without TGF-β

Cardiac fibroblasts (CFs) were isolated from the left ventricles of adult male rats. Adult male Sprague Dawley rats (200-250g) were acclimated in the facility for one week prior to isolation. Rats were given an intraperitoneal injection of 1 mL of 390 mg/mL Pentobarbitol (Fatal Plus), and after 15 minutes were checked for responsiveness before the heart was removed. The left ventricle (LV) was dissected, minced, and digested with collagenase II (1mg/mL) for 3 hours. Following centrifugation, cell pellets were resuspended and plated for 2 hours, then vigorously washed to remove non-adherent cells. The resulting cell population tested negative for VE-cadherin (endothelial cells) and positive for vimentin (fibroblasts). In all experiments, CFs were cultured on tissue culture plastic in DMEM/F12 (Gibco) supplemented with 10% FBS for a maximum of 4 days before being seeded onto hydrogels. The hydrogels were comprised of 8-arm poly(ethylene glycol) (PEG) functionalized with norbornene end groups (MW = 47 kDa), PEG thiol crosslinkers (MW = 5 kDA, JenKem), and RGD adhesive peptides. The hydrogels were photopolymerized in the presence of the photo initiator lithium phenyl-2,4,6-trimethylbenzoylphosphinate with 365 nm light for 3 minutes at an intensity of 5 mW/cm^2^. The hydrogels had a Young’s modulus of 6 kPA. Separate cohorts of CF-seeded hydrogels were treated with and without 10 ng/ml of TGF*-β* for five days before fixing and staining following the same protocol performed on VICs (see Online Methods). The CFs grown without TGF*-β* largely showed a quiescent phenotype whereas CFs grown with TGF*-β* appear activated (Supplemental Fig. 4 a&b).

#### Comparison of VIC-trained vs CF-trained models

The logistic regression models trained on the VIC dataset (Fig. 4b) were used to grade the CF dataset (Supplemental Fig. 4c). In addition, logistic regression models that were trained directly on the CF dataset were evaluated for comparison (Supplemental Fig. 4c). Interestingly some morphological features performed equivalently, regardless of being trained on the VIC or CF dataset (cell and nuclear area, 67% and 68% respectively). Other morphological features such as cell and nuclear aspect ratio performed poorly on the CF dataset when trained on the VIC dataset (33% and 37% respectively), but markedly better when trained on the CF dataset (66% and 66% respectively). In addition, the logistic regression model for the DOA variable and the MLR model performed distinctively better when trained on the CF dataset (85% and 86% respectively) compared to the VIC dataset (51% and 71% respectively). These findings suggest that the morphology of activated VICs and CFs differ significantly. However, these results demonstrate the flexibility of the models and their ability to be easily tuned to grade other fibroblast cells with high accuracy without having to repeat the Fourier-series shape analysis and PCA.

To validate the performance of the logistic regression models generated for each feature, we compared their predictions with manual phenotypic grading of the fluorescent imaging datasets comprised of CFs stained for their cytoplasm, nuclei (DAPI), and αSMA stress fibers (Supplemental Fig. 5a). Similarly to the VIC dataset (Fig. 5), k-fold cross validation (k = 10) was used to evaluate the performance of the CF trained models (Supplemental Fig. 5). The CF dataset was split into 10 random groups with 9 groups used to train the logistic regression models and the 10th serving as the validation data set (Supplemental Fig. 5b). This process was iterated 10 times so that each group served as the validation data set exactly once. The percentage of cell phenotypes that were correctly predicted in the validation data set by the logistic regression models was recorded for every iteration (Supplemental Fig. 5c). The validation showed that cell and nuclear area as well as cell and nuclear aspect ratio were good predictors of VIC phenotype with an accuracy of 67±8%, 68±7%, 66±8%, and 66±8% respectively. The DOA variable had the highest predictive capability with an accuracy of 85±3%. Beyond assessing the predictive capabilities of individual features, we also assessed the combination of all features on predicting CF phenotypes using a MLR model (Supplemental Table 7). Overall, the MLR model showed the best predictive capabilities with an accuracy of 86±3%; however, combinations of features did not result in any significant improvements in the predictions compared to the DOA variable alone.

The spectrum of CF activation was also compiled for three representative cases (Supplemental Figure 6). Interestingly, the VIC-trained MLR model seemed to consistently underestimate the activation levels of CFs compared to the CF-trained model.

**Supplemental Figure 4:**
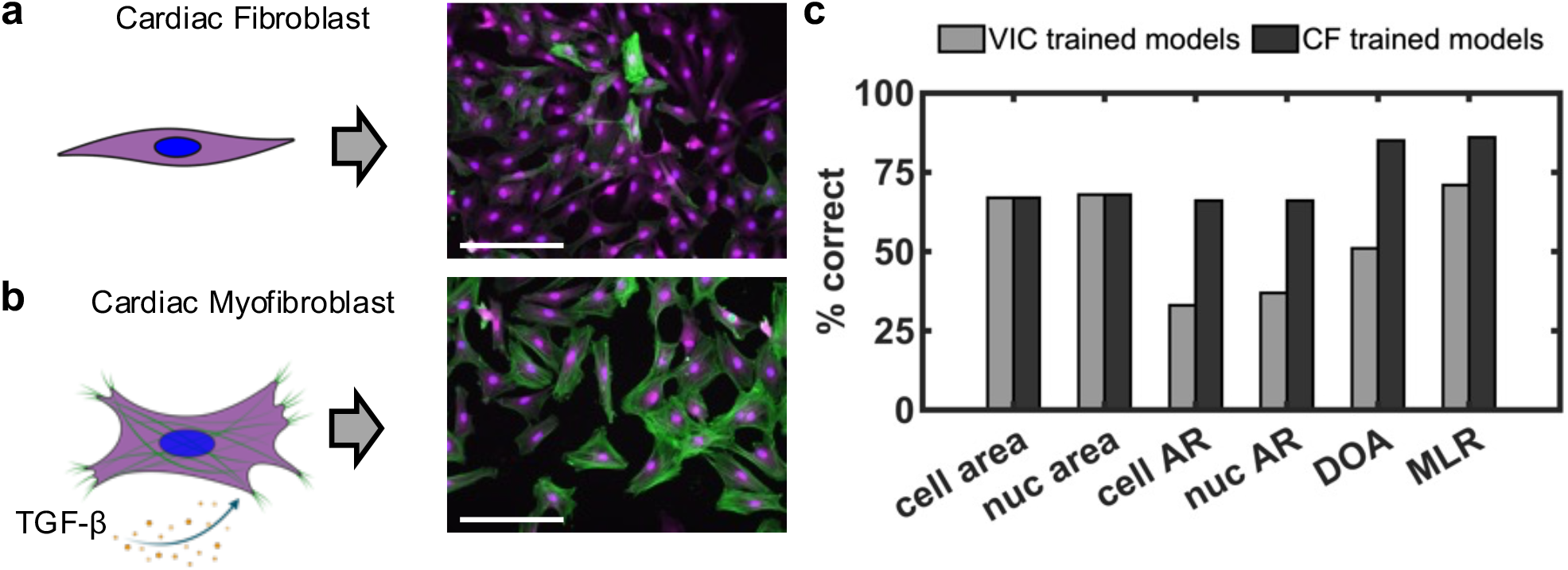
The performance of logistic regression models on an additional dataset comprised of fluorescent images of cardiac fibroblasts (CFs) cultured with and without TGF-β to induce myofibroblast activation. **a**, CFs seeded without TGF-β exhibit a quiescent phenotype characterized by limited expression of αSMA stress fibers. **b**, CFs seeded with TGF-β exhibit a transition to an activated myofibroblast phenotype characterized by high expression levels of αSMA. **c**, The percentage of CFs graded correctly for each selected feature and the MLR as compared to manual grading. Models trained on the VIC dataset were employed to grade CFs, evaluating the extent to which these models generalize beyond their training data. Grading was also performed by models trained directly on the CFs for comparison. Stained images: green - αSMA, magenta – cytoplasm, blue-nuclei. Scale bars: 200 *µ*m. Schematics created with BioRender.com.

**Supplemental Figure 5:**
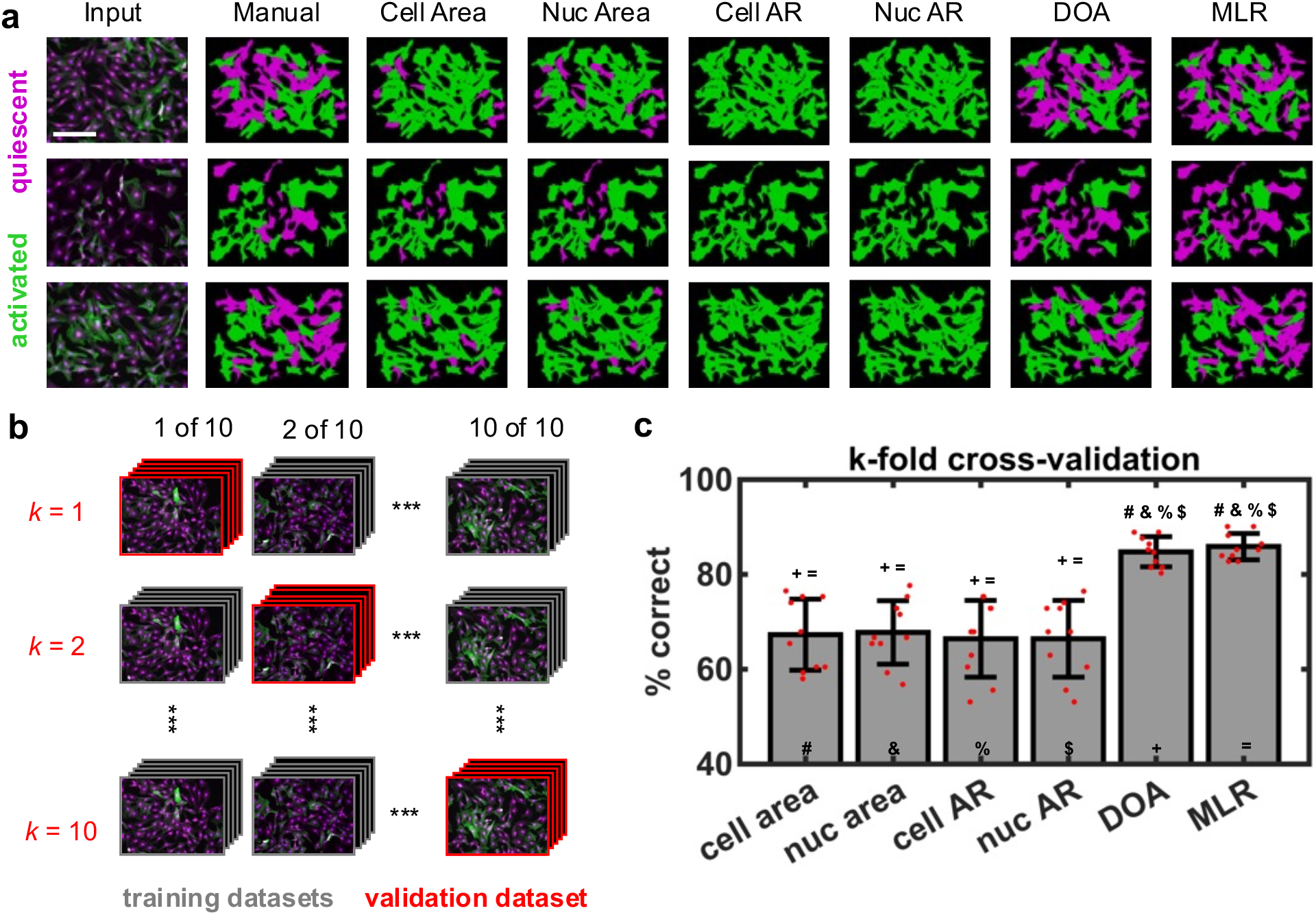
Validation of selected features toward grading activated and quiescent CFs. **a**, Three representative examples of the grading process depicting the input image, manual grading, grading done by quantitative thresholding of selected features (cell area, nuclear area, cell aspect ratio, nuclear aspect ratio, and DOA) and grading done via a multivariable logistic regression (MLR) model that combines information from all the selected features. Quantitative thresholds were determined via logistic regression. **b**, To assess the predictive capabilities of the selected features, *k*-fold cross-validation was performed in which the dataset was randomly partitioned into *k* equal sized subsets. Of the *k* subsets, *k-1* subsets were used to train logistic regression models and the remaining subset was used as the validation subset. The cross-validation process was repeated *k* times, with each of the *k* subsets serving as the validation subset exactly one time. For this analysis, *k* = 10. **c**, Bar plots showing compiled *k*-fold cross-validation results depicting the percentage of CFs graded correctly for each selected feature and the MLR when compared to manual grading. Each feature is represented with a symbol. The symbols on top of each boxplot denote the features that are significantly different from the current feature as determined by a one-way ANOVA (p < 0.01). Stained images: green - αSMA, magenta – cytoplasm, blue-nuclei. Scale bars: 200 *µ*m.

**Supplemental Table 7:** Multivariable logistic regression model coefficients for the CF dataset. n.s. – P > 0.05, *P ≤ 0.05, **P ≤ 0.01, ***P ≤ 0.001.

| Feature | Coefficient | Std. Error | P-value |
| --- | --- | --- | --- |
| Intercept ($\beta_0$) | -4.90 | 0.81 | *** |
| Cell area ($\beta_1$) | 4.55e-5 | 1.26e-4 | n.s. |
| Nuclear area ($\beta_2$) | 0.0067 | 0.004 | n.s. |
| Cell aspect ratio ($\beta_3$) | -0.68 | 0.14 | *** |
| Nuclear aspect ratio ($\beta_4$) | -0.34 | 0.53 | n.s. |
| DOA ($\beta_5$) | 14.4210 | 1.05 | *** |

**Supplemental Figure 6:**
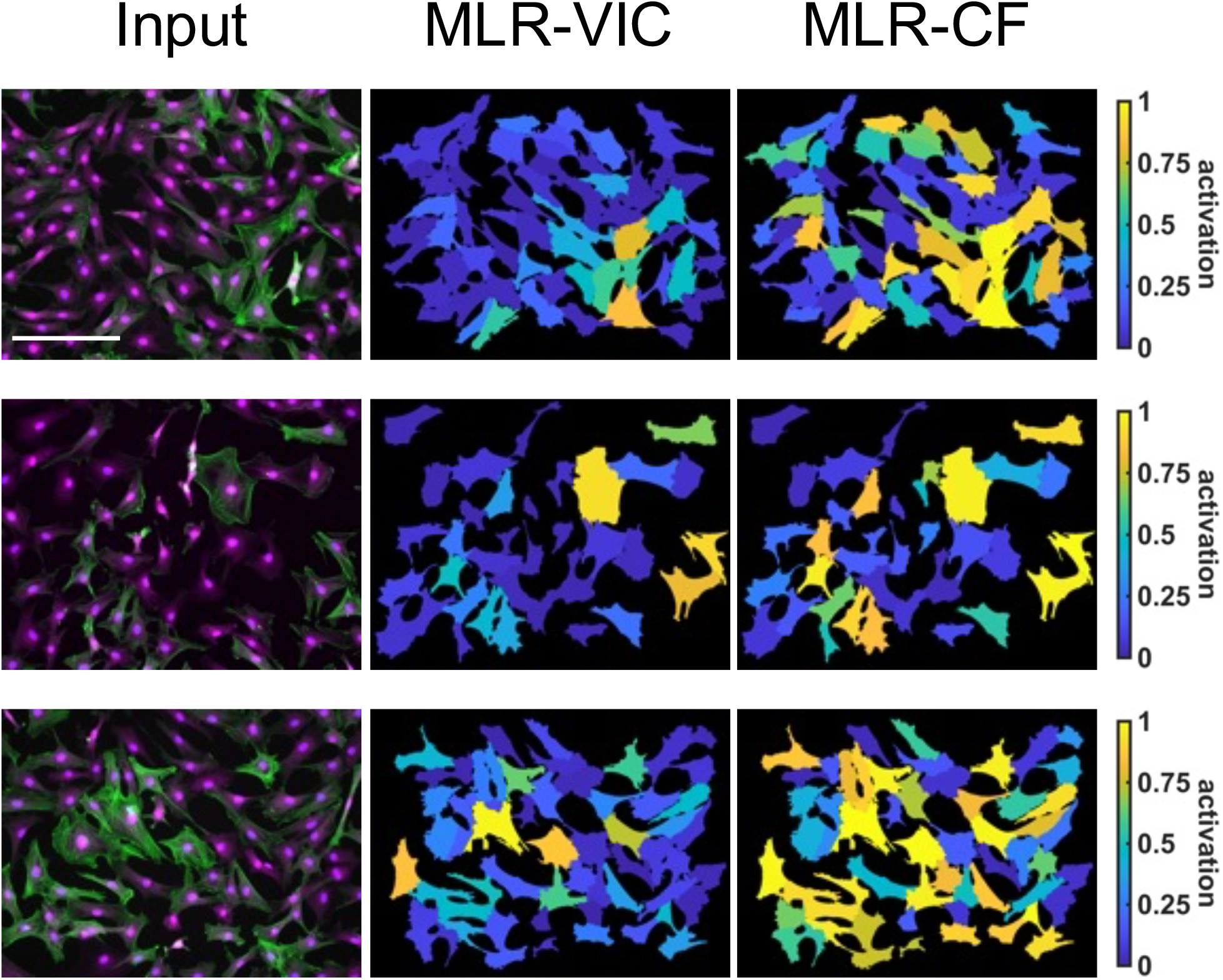
Three representative examples comparing the ability of multivariable logistic regression models trained on the VIC data set and the CF data set to grade CF activation on a spectrum. Stained images: green - αSMA, magenta – cytoplasm, blue-nuclei. Scale bars: 200 *µ*m.

